# *SORBS2* is a susceptibility gene to arrhythmogenic right ventricular cardiomyopathy

**DOI:** 10.1101/725077

**Authors:** Yonghe Ding, Jingchun Yang, Peng Chen, Tong Lu, Kunli Jiao, David Tester, Kai Jiang, Michael J Ackerman, Yigang Li, Dao Wu Wang, Dao Wen Wang, Hon-Chi Lee, Xiaolei Xu

## Abstract

**BACKGROUND:** Arrhythogenic right ventricular cardiomyopathy (ARVC) is an inherited cardiomyopathy characterized by right ventricular remodeling and ventricular arrhythmia. To date, 16 ARVC causative genes have been identified from human genetic studies, accounting for about 60% of ARVC probands. Genetic basis for the remaining 40% ARVC probands remain elusive.

**METHODS:** Prompted by a zebrafish mutagenesis screen that suggested the *Sorbin and SH3 domain-containing 2 (SORBS2)* ortholog as a candidate cardiomyopathy gene, we conducted detailed expressionl analysis of Sorbs2 in mice, as well as phenotypic characterization in the Sorbs2 knock-out (KO) mice. The intercalated disc (ICD) expression pattern and ARVC-like phenotypes further prompted us to conduct targeted sequencing of human patients with ARVC to search for rare variants in the *SORBS2* gene.

**RESULTS:** *Sorbs2* is robustly expressed in the mouse heart, encoding an adhesion junction/desmosome protein that is mainly localized to the ICD. A mutation with near complete depletion of the Sorbs2 protein in mouse results in phenotypes characteristic of human ARVC, such as dilated right ventricle (RV), RV dysfunction, spontaneous ventricular tachycardia (VT), and premature death. Sorbs2 is required to maintain the structural integrity of ICD. Its absence resulted in profound cardiac electrical remodeling with impaired impulse conduction and action potential derangements. Five rare variants were identified from a cohort of 59 ARVC patients, among which two variants affect splicing.

**CONCLUSIONS:** Sorbs2 KO mouse is an ARVC model and *SORBS2* is a new ARVC susceptibility gene.

## INTRODUCTION

Arrhythmogenic right ventricular cardiomyopathy (ARVC), also referred to as arrhythmogenic right ventricular dysplasia (ARVD), is a rare inherited cardiac muscle disorder that can lead to heart failure and sudden cardiac death (SCD)^1^. The prevalence of ARVC in the general population is between 1:1,000 and 1:5,000, accounting for up to 50% SCD in the United States^2-4^. ARVC is characterized by structural abnormalities in the right ventricle (RV) and ventricular arrhythmia at early stages. Left ventricular (LV) involvement becomes common with disease progression, and the biventricular involvement prompted recent recommendation of using a broader term, arrhythmogenic ventricular cardiomyopathy (ACM), for this disease^5, 6^. Current management of ARVC is mostly palliative and focuses on delaying disease progression and preventing SCD. No curative treatment is available for this life-threatening disease^1^.

Human genetic studies of ARVC probands have linked 16 genes to ARVC to date^1, 7^, among which seven encode desmosomal proteins located in the intercalated disc (ICD). Mutations in desmosomal protein coding genes, such as *PKP2, DSP* and *DSG2*, account for up to 63% of the ARVC probands^1, 8^. A small percentage of ARVC probands have sequence variants in non-desmosomal protein-encoding genes such as *CTNNA3* and *CDH2*^*9*, *10*^. *Mouse models for five desmosome/ICD protein coding genes (DSP, DSG2, DSC2, PKP2, JUP*) have been generated through either targeted genetic deletion or transgenic overexpression of genes with disease-causing mutations^8, 11^. The key features of human ARVC phenotypes such as structural abnormalities in the ventricles, spontaneous ventricular tachicardia (VT), and SCD, can be recapitulated in the majority of these mouse models. Certain aspects of human ARVC, such as fibrofatty infiltration, however, was only noted in the mouse model involving *DSP*^12, 13^, *but not those involving DSG2, DSC2* and *JUP* ^*8*, *14*^. *Because of the convenience of embryonic morpholino knock-down and amenability to high-throughput genetic and compound screening, zebrafish ARVC models for DSC2* and *JUP* have also been developed^15, 16^.

Through an insertional mutagenesis screen in zebrafish, we recently identified the gene-breaking transposon *(GBT) 002* as a mutant that exerted deleterious modifying effects on doxorubicin-induced cardiomyopathy (DIC) ^17^. *GBT002* was later determined as a loss-of-function mutant that disrupts *sorbs2b (sorbin and SH3 domain containing 2b)*, one of the two *SORBS2* orthologues in zebrafish ^17^. *SORBS2*, also known as Arg/c-ab1 kinase binding protein 2 (*ArgBP2*), encodes a member of the SOHO adaptor family protein that contains a Sorbin domain at its N terminus and three SH3 domain at its C terminus ^18^. The encoded protein manifests a restricted subcellular expression pattern, including association with stress fibers ^19^, and sarcomeric Z discs in cardiomyocytes ^20, 21^. Besides our evidence from zebrafish, several recent studies also suggested the cardiac involvement of Sorbs2. First, knock-down of *Sorbs2* in neonatal rat cardiomyocytes in culture resulted in cellular hypertrophy ^22^. Second, *SORBS2* was a candidate gene in regions of copy number variations linked to congenital heart disease in human ^23^. Third, expression of SORBS2 protein was found to be significantly increased in the sera of patients with acute myocardial infarction, but depleted in the infarcted myocardia ^24^. However, *in vivo* cardiac phenotypes have not been reported in a *Sorbs2* knock-out (KO) mouse model ^25^.

Here, we reported detailed cardiac studies of Sorbs2 in mouse. We revealed that Sorbs2 encodes an ICD protein, and *Sorbs2* KO mice exhibit ARVC-like phenotypes including right ventricular cardiomyopathy, VT, and premature death. Scanning a cohort of 59 Chinese Han patients with ARVC led to the identification of five rare sequence variants. Collectively, our results support that the Sorbs2 KO mouse is an ARVC model and *SORBS2* as a previously unrecognized susceptibility gene to ARVC.

## METHODS

An expanded Materials and Methods is provided in the online-only Data Supplement. All the supporting data and materials within the article will be made available upon reasonable request.

### Statistical Analysis

Kaplan-Meier survival curve, echocardiography (ECG) and magnetic resonance imaging (MRI) analyses are from cumulative data. The unpaired two-tailed Student’s *t*-test was used to compare 2 groups. One- or two-way analysis of variance (ANOVA) was used to assess differences among multiple groups, as appropriate. Tukey test was applied for the post hoc analysis to confirm the diiference. The log-rank test was used to determine the difference in animal survival. Chi-square test was used to determine the enrichment of pathogenic variants in *SORBS2* gene from ARVC patients. All quantitative data are presented as mean ± standard deviation (SD). *P* values less than 0.05 were considered to be significant. All statistical analyses were performed with GraphPad Prism 7 software.

## RESULTS

### Sorbs2 is an intercalated disc protein in cardiomyocytes

In mouse, there is a single *Sorbs2* gene in chromosome 8 (NM_001205219), encoding a protein that is highly conserved with human SORBS2, especially in its N-terminal sorbin homologue domain and C-terminal SH3 domain where they share more than 95% homology (Figure 1 in the online-only Data Supplement). Expression of Sorbs2 protein is highly enriched in the mouse heart (Figure 1A). The mouse cardiac Sorbs2 is around 120 kilo Dalton (kDa) in size, whereas its brain isoform is about 150 kDa (Figure 2 in the online-only Data Supplement) ^25^, suggesting there exists tissue-specific alternative splicing events and/or post-translational modification. At the subcellular level, Sorbs2 is localized strongly in the intercalated discs (ICD) of cardiomyocytes, with weaker localization in the perinuclear areas and the Z discs (Figure 1B). To resolve the Sorbs2 expression within ICD, we conducted double immunostaining using a Sorbs2 antibody and antibodies against β-catenin, plakoglobin (Pkg), or connexin 43 (Cx43), markers of the adhesion junction, desmosome, or gap junction structures, respectively. We found that Sorbs2 mostly co-colocalizes with β-catenin, partially with Pkg, while only slightly with Cx43 (Figure 1C). The weighted co-localization efficiency of Sorbs2 is about 60% with β-catenin, 20% with Pkg, and 6% with Cx43, suggesting that Sorbs2 is mainly localized in the adhesion junction, and partially in the desmosome.

**Figure 1.**
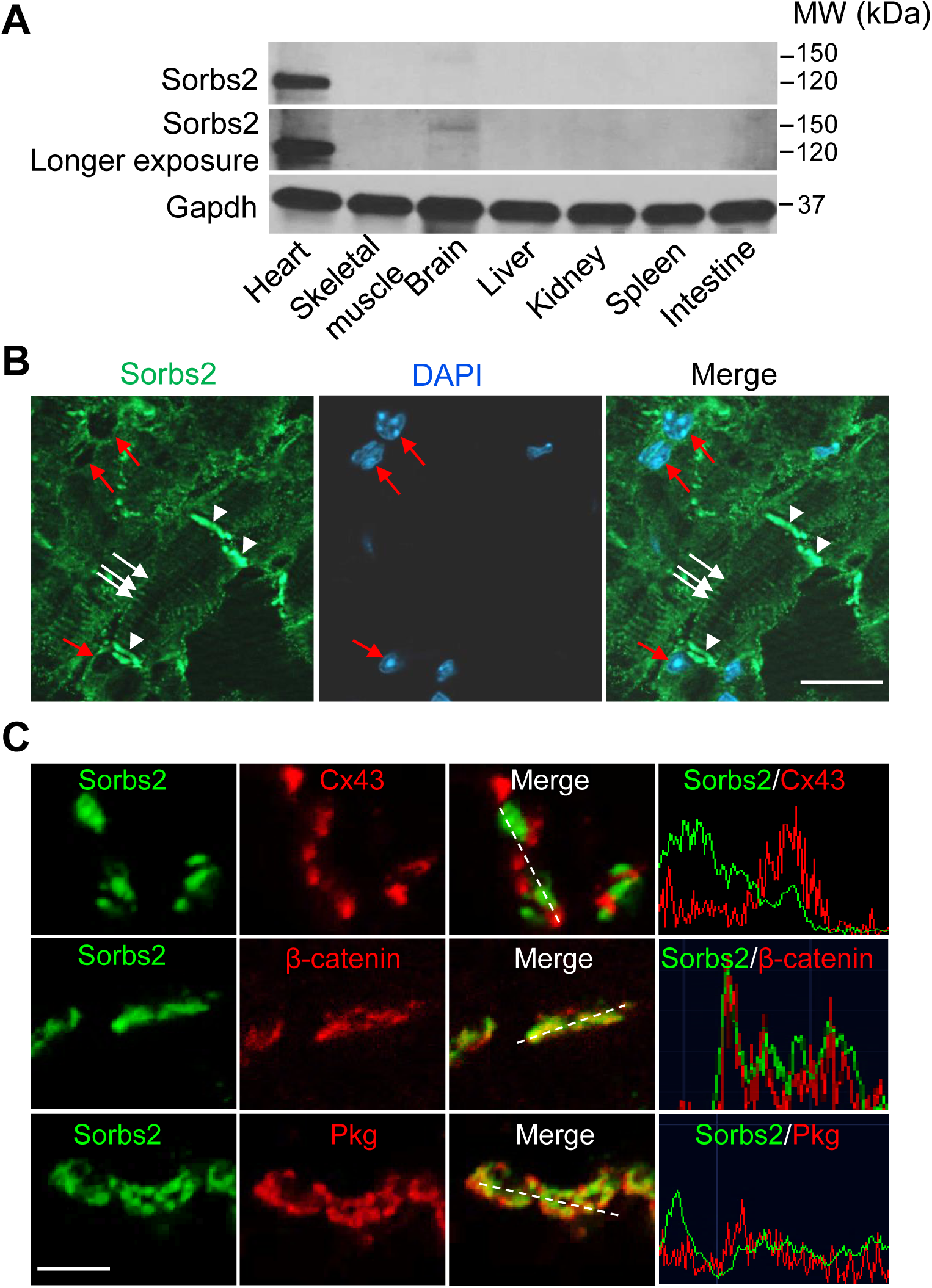
Sorbs2 is a cardiac enriched intercalated disc and desmosome protein in mouse. **A**, Western-blot analysis of the Sorbs2 protein expression in different mice tissues. **B**, Images of immunostaining using an anti-Sorbs2 antibody and counter stained with 4′,6-diamidino-2-phenylindole (DAPI) in sectioned WT mouse hearts. Arrowheads point to intercalated disc (ICD). While arrows point to Z-disc localization. Red arrows point to perinuclear. Scale bar, 20 µm. **C**, Shown are immunostaining and pseudo-line analysis of sectioned mouse hearts, indicating co-localization of Sorbs2 protein mostly with β-Catenin and Plakoglobin (Pkg), but less with Connexin 43 (Cx43). Scale bars: 2 µm.

**Figure 2.**
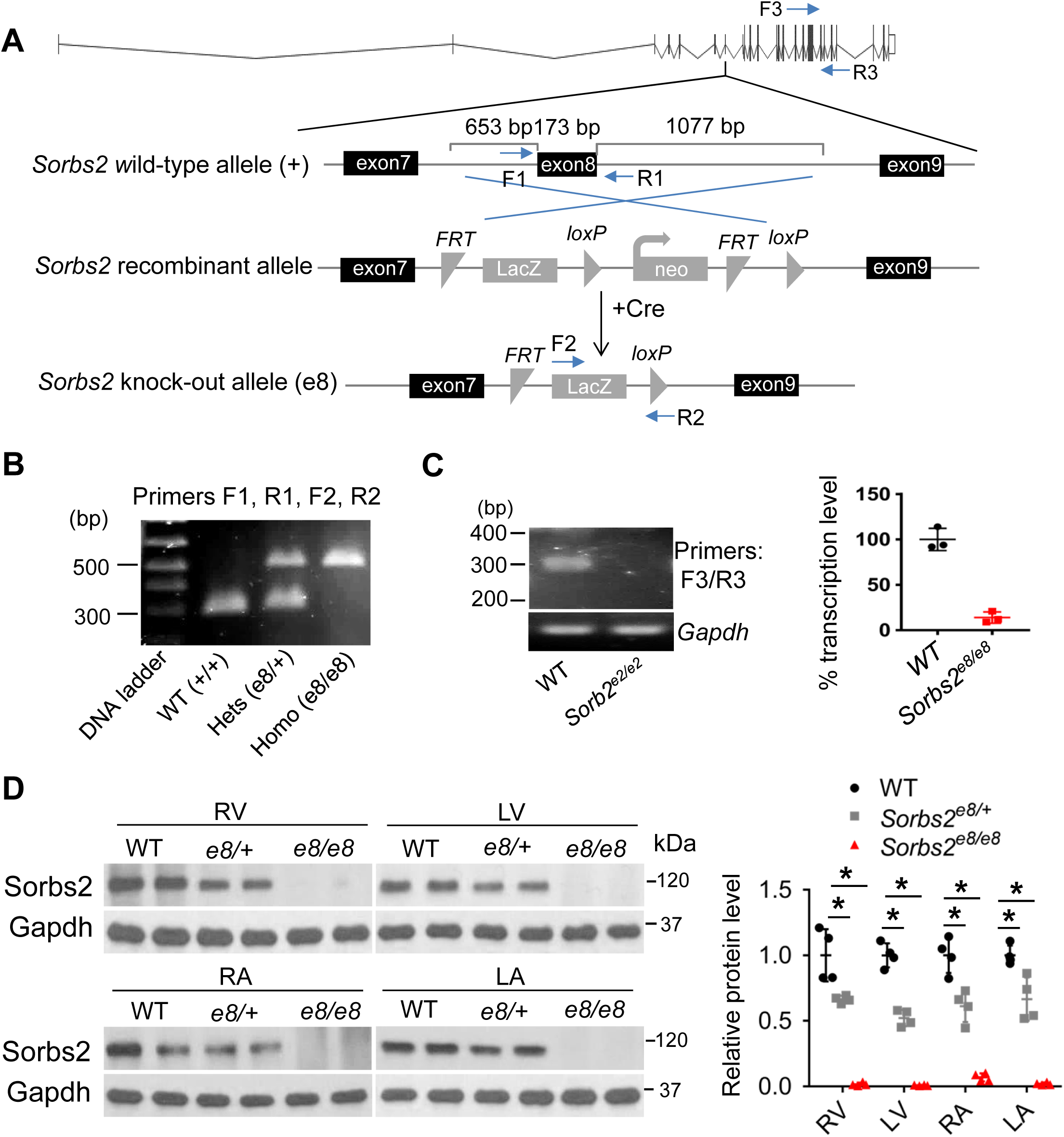
Targeted deletion of exon 8 in *Sorbs2* gene led to near null depletion of the whole Sorbs2 transcript and protein in mouse. **A**, Schematic illustration of targeted deletion of Sorbs2 Exon 8 in the *Sorbs2*^*e8/e8*^ mice and locations of primers used for genotyping PCR and transcription analysis. **B**, Rrepresentative DNA gel images of PCR genotyping for identifying WT (292 bp), *Sorbs2*^*e8/+*^ heterozygous (hets), and *Sorbs2*^*e8/e8*^ homozygous (homo) mutant alleles (500 bp). **C**, Representative DNA gel images and quantitative RT-PCR analysis of the Sorbs2 transcripts in WT and *Sorbs2*^*e8/e8*^ mutants. **D**, Western-blot and quantification of Sorbs2 protein expression in each cardiac chamber of WT and *Sorbs2*^*e8/e8*^ mutants. N=4. Two-way ANOVA. *, p<0.05. RV, right ventricle. LV, left ventricle. RA, right atrium. LA, left atrium.

### Deleterious modifying effects of *GBT002/Sorbs2* on DIC is cardiomyocyte-primary and conserved in mouse

To elucidate cardiac functions of Sorbs2, we first studied the doxorubicin-induced cardiomyopathy (DIC) modifying effects in the zebrafish gene-breaking transposon (*GBT)002* mutant. *Because* the two Loxp sites in the GBT vector enables cardiomyocyte-specific genetic manipulation ^26^, we crossed *Tg(cmlc2:cre-ER)*, a myocardium-specific expressed and hydroxytamoxifen inducible Cre transgenic fish line ^27^, into the *GBT002/sorbs2b* muant for a cardiomyocyte-specific rescue experiment. We found that the mutagenic insertion was effectively removed and *sorbs2b* transcript was fully restored in the myocardium of the *GBT002*^*-/*-^ *;Tg(cmlc2:cre-ER)* double mutant/transgenic fish upon hydroxytamoxifen treatment (Figure 3 in the online-only Data Supplement). While *GBT002*^*-/*-^ homozygous mutant manifests no visually noticeable phenotypes, after doxorubicin (DOX) stress, the *GBT002*^*-/*-^ mutant exhibited significantly deleterious DIC modifying effect, as indicated by the increased fish mortality (Figure 2D in the online-only Data Supplement). This deleterious DIC modifying effect of *GBT002/sorbs2b* was largely rescued in the *GBT002*^*-/*-^*;Tg(cmlc2:cre-ER)* double mutant/transgenic fish, suggesting a critical contribution of myocardial expression of *sorbs2b* to its DIC modifying effects.

**Figure 3.**
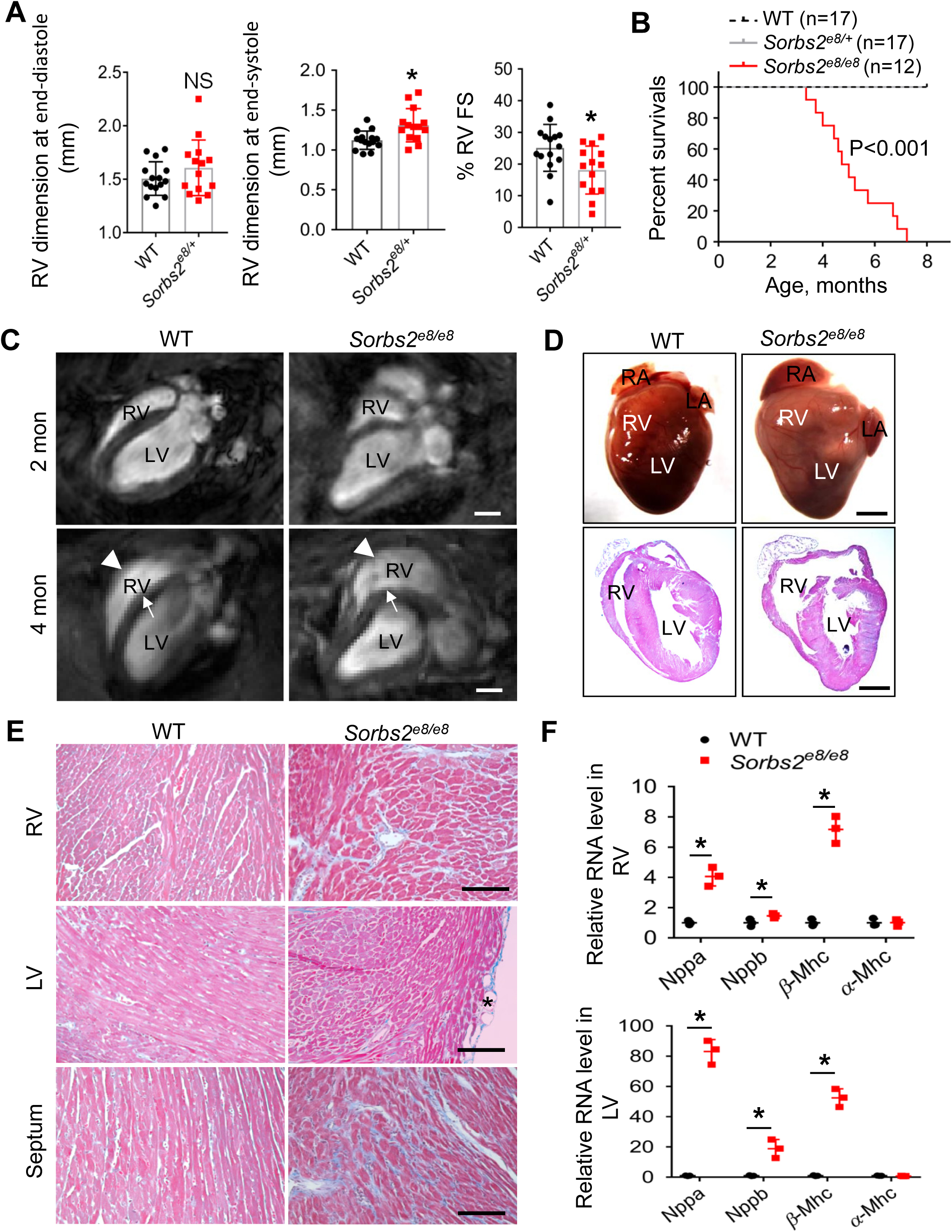
Sorbs2 deficiency leads to RV dilation, cardiac dysfunction and premature death. **A**, Echocardiography indices in the right ventricle (RV) of *sorbs2*^*e8/*-^ and WT control mice at 4 weeks post-doxorubicin injection (12 mg/kg body weight). N=14-15. Two-paired student’s *t*-test. FS, fractional shortening. **B**, Kaplan-Meier survival curves of *Sorbs2*^*e8/e8*^ and WT control mice. N=12-17. Log-rank test. **C**, Long-axis views of magnetic resonance imaging (MRI) images at 2 and 4 month-old mice. RV dilation (arrow heads) is prominent in the *Sorbs2*^*e8/e8*^ mouse hearts at 4 months old. Note that septum bending defect (arrows) was also detected. **D**, Images of dissected whole hearts (upper) and cross-sectional view of Hematoxylin and eosin (H&E) stained hearts (lower). Dramatically enlarged RV and protuberant were noted in the *Sorbs2*^*e8/e8*^ mice. Note that both RA and LA enlargement was also frequently observed. **E**, Representative images of masson’s trichrome staining demonstrated myocardium fibrosis in the RV and septum, but not in the LV of *Sorbs2*^*e8/e8*^ mice. Scale bar: 50 μm. **F**, Quantitative RT-PCR analysis of heart failure markers in RV and LV of the *Sorbs2*^*e8/e8*^ mice hearts compared to WT. RV, right ventricle. LV, left ventricle. n=3, *: p<0.05. Two-way ANOVA.

To prove the conservation of the DIC modifying effects in mammals, we obtained the *Sorbs2*^*e8*^ knock-out (KO) mice (B6N(Cg)-Sorbs2 /J), in which a targeted deletion of 1903 bp nucleotides removes part of intron 7, the whole exon 8, and part of intron 8 from the *Sorbs2* genomic locus (Figure 2A, 2B). The exon 8 encodes part of the N-terminal Sorbin homologue domain (Figure 1 in the online-only Data Supplement), removal of which results in aberrant splicing of exon 7 to an ectopic exon encoding LacZ (Figure 2A, 2B and Figure 3D, 3E in the online-only Data Supplement), leading to dramatical reduction of the whole Sorbs2 transcript (Figure 2C). At the protein level, the Sorbs2 protein is about 50% reduced in the *Sorbs2*^*e8/+*^ heterozygous and nearly completely absent in the *Sorbs2*^*e8/e8*^ homozygous mice in all 4 cardiac chambers, as indicated by Western blots using an antibody that recognizes a C-terminal antigenic epitope of the Sorbs2 protein (Figure 2D, and Figure 1 in the online-only Data Supplement). In the *Sorbs2*^*e8/e8*^ mice, Sorbs2 protein in the muscle and its larger isoform in the brain are also mostly ablated (Figure 2A in the online-only Data Supplement).

Consistent with the zebrafish data, we noted deleterious modifying effects of *Sorbs2*^*e8/+*^ heterozygous mice on DIC. Mild but significant reduction of right ventriclular (RV) function was noted upon DOX stress due to enlarged end-systolic diameter (Figure 3A). Other cardiac pump indice remained unchanged (Table 1 in the online-only Data Supplement). We did not conduct detailed studies of the DIC modifying effects of *Sorbs2*^*e8/e8*^ homozygous mice, because of the interference of severe cardiac phenotypes, as will be decribed below.

**Table 1.** Quantification of cardiac function indices in *Sorbs2* ^*e8/e8*^ mutant and WT control mice via magnetic resonance imaging (MRI) at 4 months old.

|  | WT siblings | *Sorbs2<sup>es8/es8</sup>* |
| --- | --- | --- |
| Mice number (n) | 5 | 9 |
| Body weight (g) | 24.66 ± 1.05 | 21.44 ± 1.33* |
| Heart weight/body weight (mg/g) | 6.41 ± 0.61 | 9.24 ± 0.78* |
| RVEDV (μL) | 34.12 ± 10.45 | 37.22 ± 11.32 |
| RVESV (μL) | 13.38 ± 4.19 | 21.93 ± 7.3* |
| RVEF(%) | 58.69 ± 12.68 | 41.17 ± 9.9* |
| RVFACI (%) | 46.79 ± 6.08 | 32.37 ± 5.63* |
| RVFACs (%) | 57.12 ± 15.73 | 29.92 ± 14.85* |
| LVEDV (μL) | 46.23 ± 9.09 | 31.38 ± 7.67* |
| LVESV (μL) | 15.76 ± 6.4 | 16.12 ± 7.2 |
| LVEF (%) | 66 ± 9.05 | 50 ± 16.11* |
| IVSs (mm) | 1.35 ± 0.07 | 1.09 ± 0.07* |
| IVSd (mm) | 0.95 ± 0.05 | 0.84 ± 0.05 |
| LVPWs (mm) | 2.2 ± 0.11 | 2.12 ± 0.11 |
| LVPWd (mm) | 1.14 ± 0.11 | 1.55 ± 0.11* |

RVEDV: right ventricle end-diastolic volume; RVESV: right ventricle end-systolic volume; RVEF: right ventricle ejection fraction; RVFACI: right ventricle fractional area change along long-axis. RVFACs: right ventricle fractional area change along short-axis. LVEDV: left ventricle end-diastolic volume; LVESV: left ventricle end-systolic volume; LVEF: left ventricle ejection fraction; IVSs, Interventricular septum thickness at end-systole; IVSd, Interventricular septum thickness at end-diastole; LVPWs, Left ventricular posterior wall thickness at end-diastole; LVPWd, left ventricular internal dimension at end-diastole. \*, $P < 0.05$ , unpaired student *t* test.

### Depletion of Sorbs2 leads to RV dilation, cardiac dysfunction, premature death, and ventricular arrhythmia in mice

*Sorbs2*^*e8/e8*^ homozygous mice are fertile, but started to die at about 3.5 months old, and in rare cases they could survive to 7 months (Figure 3B). Using the non-invasive cardiac magnetic resonance imaging (MRI), we detected prominent RV dilation in the *Sorbs2*^*e8/e8*^ mice as early as 2 months and it got worse at 4 months (Figure 3C). The RV bulges out along the short axis, and the overall shape of the ventricle changes from crescent to triangle-like in lateral view (Figure 3D). The ventricular septum in the *Sorbs2*^*e8/e8*^ mice becomes progressively bended, ranging from mild at 2 months to severe at 4 months, in contrast to the straight ventricular septum in wild type (WT). Both heart weight and left ventricular posterior wall thickness at end diastole (LVPWd) are increased significantly, suggesting the presence of LV hypertrophy (Table 1). Cardiac function in both RV and LV drop significantly. In association with RV dilation at 4 months, fibrosis was detected in both RV and LV, and more prominently in the septum (Figure 3E). Occasionally, adipose tissue deposition was noted in the LV, but not in other chambers. Transcription levels of molecular markers for heart failure such as *Nppa, Nppb* and *β-Mhc* are markedly elevated in both LV and RV (Figure 3F).

Besides the profound cardiac structural remodeling, abnormal electrical remodeling was also striking in the *Sorbs2*^*e8/e8*^ mice. Significant increase in QRS and S wave durations with the development of right bundle branch block (RBBB) are commonly detected in the *Sorbs2*^*e8/e8*^ mice at 4 months of age (Figure 4A). Random 2-minute electrocardiogram (ECG) recordings in anesthetized animals showed spontaneous arrhythmias including premature ventricular contractions (PVCs), nonsustained ventricular tachycardias (VTs), and polymorphic VTs in the *Sorbs2*^*e8/e8*^ mice (7/31), but not in WT controls (0/37) (Figure 4B) (χ^2^, *p*<0.05, Chi-square test). Electrophysiological studies in Langendorff-perfused hearts showed that the ventricular effective refractory period (VERP) is markedly prolonged in *Sorbs2*^*e8/e8*^ mice compared to WT (Figure 4C). Sustained VTs with durations greater than 30 seconds were induced by programmed electrical stimulation or burst pacing in 9 out of 10 *Sorbs2*^*e8/e8*^ hearts (90%), but only in 3 out of 12 WT hearts (25%) (χ^2^, *p*<0.05, Chi-square test). These results suggest that the *Sorbs2*^*e8/e8*^ mice have a greater propensity to develop spontaneous and induced ventricular tachyarrhythmias which might be pertinent to the shortened life-span of these animals. Intracellular recordings of action potentials from the endocardial side of RV showed marked electrophysiological abnormalities in *Sorbs2*^*e8/e8*^ mice (Figure 4D). The resting potentials are significantly depolarized in *Sorbs2*^*e8/e8*^ mice compared to WT. Action potential amplitudes and upstroke velocities (dV/dt) are significantly reduced in *Sorbs2*^*e8/e8*^ mice (Figure 4E). There are remarkable prologation of action potential durations at 50 % and 90% repolarization (APD50 and APD90), as well as shortening in the effective refractory period (ERP) detected in the *Sorbs2*^*e8/e8*^ mice.

**Figure 4.**
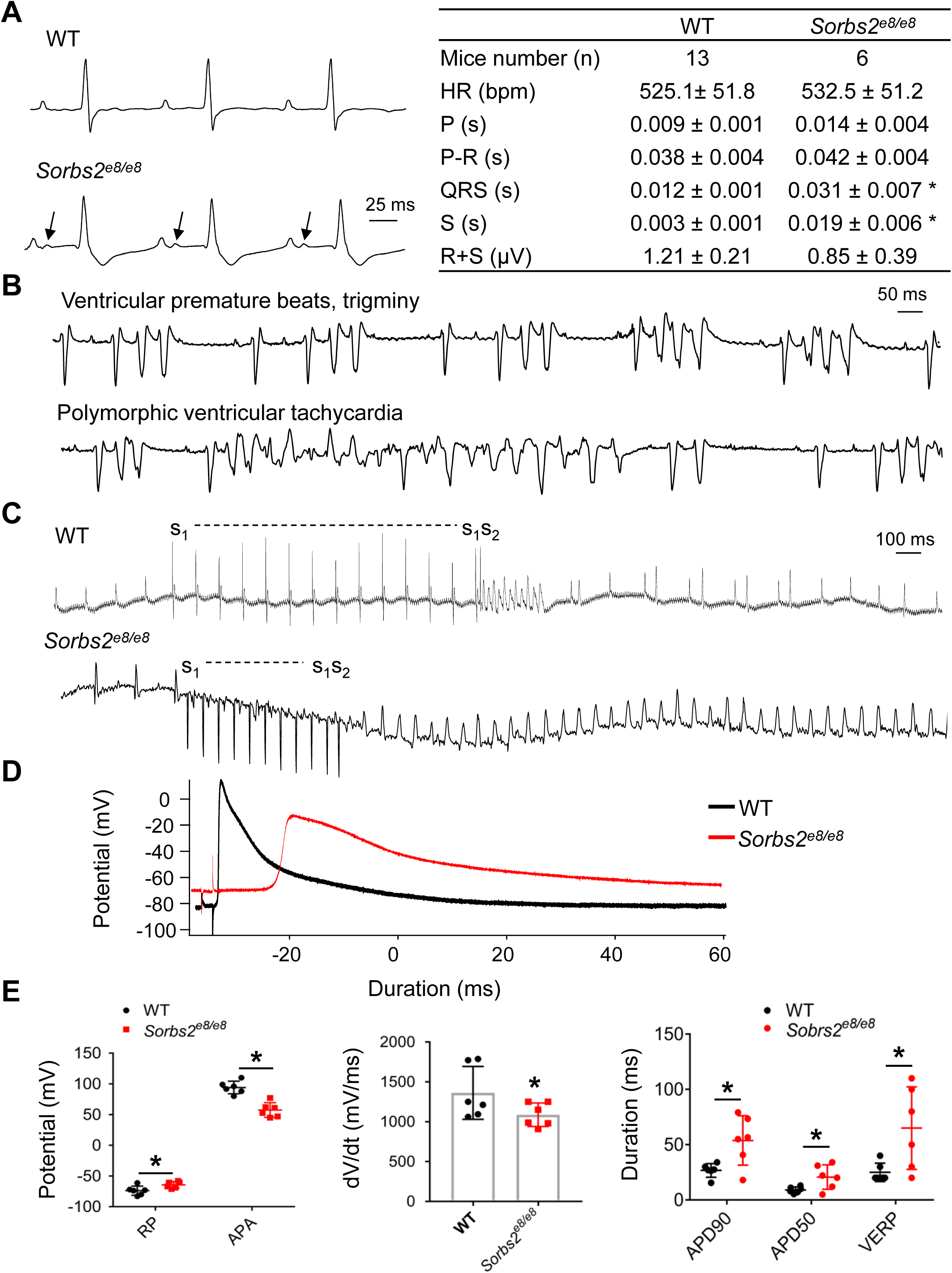
Sorbs2 deficiency leads to cardiac arrhythmia. **A**, Shown are surface electrocardiograms (ECG) and quantification analysis in 4 months old WT and *Sorbs2*^*e8/e8*^ mice. HR: heart rate; Bpm, beats per minute; P: duration of P waves; P-R: duration of PR interval; QRS: duration of QRS complex; S: duration of S wave; R+S: QRS amplitude. N=6-13. Two-paired student *t*-test. *: p<0.05. **B**, Ventricular premature beats and trigeminy (upper panel) and polymorphic ventricular tachycardia (VT) (lower panels) were recorded in *Sorbs2*^*e8/e8*^ mice. **C**, Induction of non-sustained VT in Langendorff-perfused WT mouse heart by programmed electrical stimulation (s1-s1=100 ms, s1-s2=30 ms) (upper panel); induction of sustained monomorphic VTs in Langendorff-perfused *Sorbs2*^*e8/e8*^ mouse heart by programmed stimulation (s1-s1=100 ms, s1-s2=80 ms) (lower panel). **D**, Representative action potentials were recorded from the endocardial surface of isolated RVs of *Sorbs2*^*e8/e8*^ and WT mice at a pacing cycle length of 200 ms. **E**, Compared to WT mice, *Sorbs2*^*e8/e8*^ mice RV action potentials have depolarized resting potentials (RP), decreased amplitudes and reduced upstroke velocity at phase 0 (Vmax), and prolonged action potential durations at 50% (APD50) and 90% (APD90) repolarization, as well as lengthened ventricular effective refractory period (VERP). N=6. Two-way ANOVA. *: p<0.05.

### *Sorbs2*^*e8/e8*^ manifests disrupted ICD structure and reduced expression of Cx43 protein

Prompted by the subcellular expression of Sorbs2 in ICD, we sought to examine whether Sorbs2 depletion affects the expression of other ICD proteins. Through immunostaining of sectioned hearts, we firstly confirmed depletion of the Sorbs2 expression in both ICD and Z disc in the *Sorbs2*^*e8/e8*^ hearts at 4 months of age (Figure 4 in the online-only Data Supplement). Loss of Sorbs2 does not seem to affect the protein expression levels of adhesion junction or desmosomal proteins such as β-catenin and Pkg (Figure 4 in the online-only Data Supplement); however, the expression level of the gap function protein Cx4 is markedly reduced. While the expression level of N-cadherin remains largely unchanged, its dotted expression pattern mostly seen in WT heart was changed to a continuous line in *Sorbs2*^*e8/e8*^ mutant hearts (Figure 5A), and from an alternating pattern with Pkg to a largely overlapping pattern (Figure 5 in the online-only Data Supplement). The significant reduction of Cx43, but not other ICD proteins, was confirmed using Western blot analysis (Figure 5B). Downregulation of Cx43 is known to impair electrical impulse conduction and create substrates for cardiac arrhythmias ^28, 29^. A recent study reported Sorbs2 as a potent RNA binding protein to stabilize tumor-suppressor gene expression in ovarian cancer ^30^. We thus carried out targeted RNA immuoprepicipitation (RIP) assay and found that Sorbs2 protein could bind the Cx43 mRNA (Figure 5C). Together with the data that Cx43 transcript was significantly decreased in the *Sorbs2*^*e8/e8*^ mice hearts as well (Figure 5D), we speculated that disrupted transcriptional regulation of Cx43 by Sorbs2 might contribute to the cardiac arrhythmia in the *Sorbs2*^*e8/e8*^ mice. More detailed mechanistic study along this direction may warrant further investigation.

**Figure 5.**
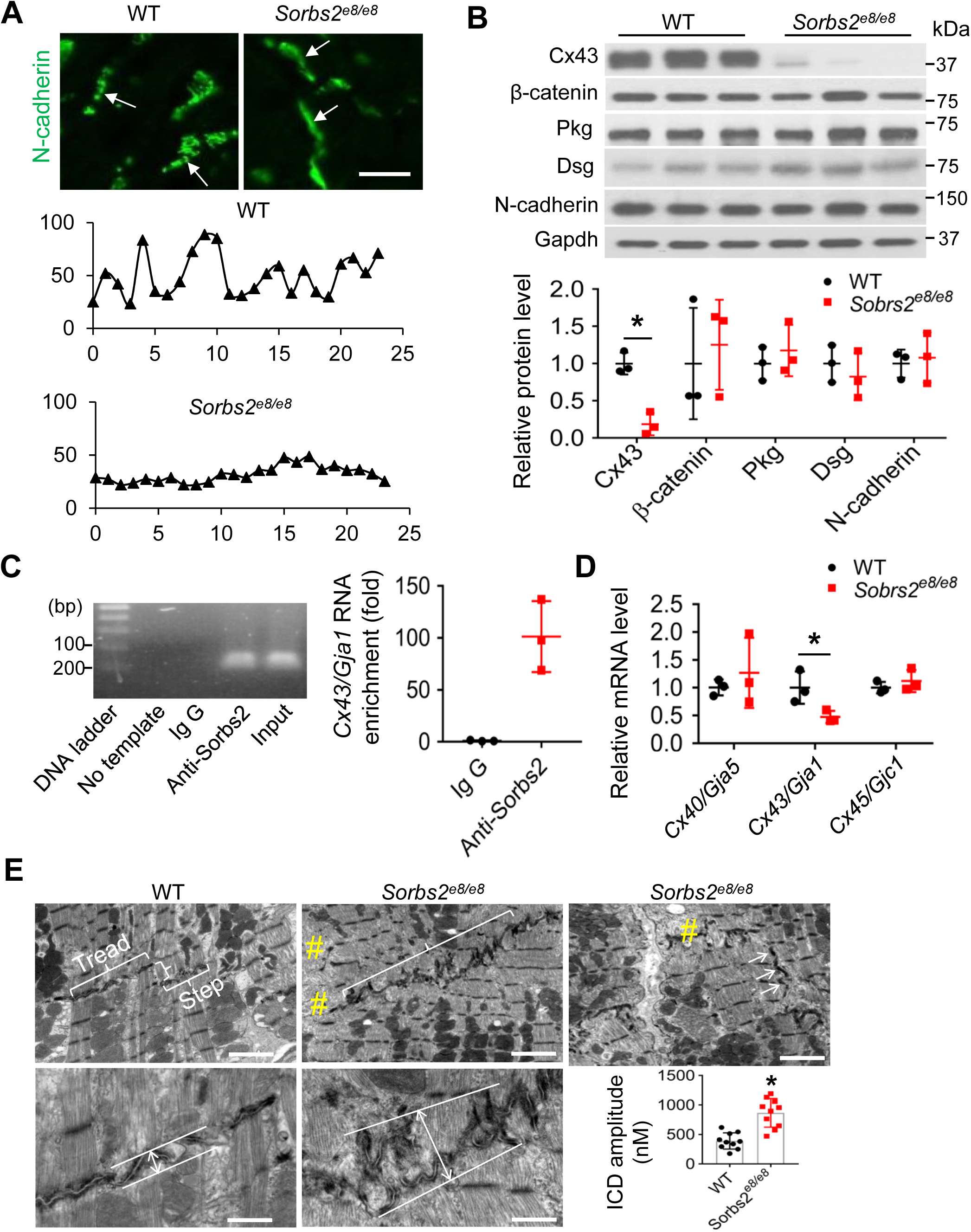
Dual function of Sorbs2 in maintaining the ICD integrity and regulation the expression of gap junction protein Cx43. **A**, Shown are ICD images and represented pseudo line analysis of mouse hearts after immunostaining using an anti-N-cadherin antibody. The expression of N-cadherin in ICD manifests as discontinuous dots in WT mice, which swithed to continuous lines in the S*orbs2*^*e8/e8*^ mice (arrows). Scale bar: 20 μm. The intensity distribution of N-cadherin signal along ICD appear as wavy lines in WT mice which largely changed to flat lines in the S*orbs2*^*e8/e8*^ mice. **B**, Western-blot and quantification of ICD proteins in WT and *Sorbs2*^*e8/e8*^ KO mice hearts at 4 months old. N=3. Two-way ANOVA. *: p<0.05. **C**, A representative DNA gel and quantitative RT-PCR analysis of the *Cx43*/*Gja1* mRNA level in the anti-Sorbs2 antibody and Ig G control immunoprecipated tissue lysates from RV of WT mice. **D**, Quantitative RT-PCR analysis of mRNA levels of indicated genes in the RV tissue of *Sorbs2*^*e8/e8*^ mice compared to WT controls. N=3. Two-way ANOVA. *: p<0.05. **E**, TEM images and schematic illustration on measuring and quantifying the ICD amplitude in WT and *Sorbs2*^*e8/e8*^ KO mice hearts at 4 months old. #: disrupted myofibril. Scale bars for upper panels, 2 μm; Scale bars for lower panels, 500 nM. *: p<0.05. Unpaired student’s *t*-test.

At the ultrastructural level, ICD in a WT mouse heart is a unique structure at the longitudinal junction between two neighboring cardiomyocytes consists of two regions: the tread region that is typically in parallel with Z discs along the myofibrils, and the step region that connects the tread regions and is typically perpendicular to the tread and Z discs (Figure 5E). This highly organized ICD structure connects the myofibrils from two neighboring cardiomyocytes across the membrane, ensuring coordinated contraction ^31^. In the *Sorbs2*^*e8/e8*^ mice hearts at 4 months old, however, there is no clear separation between the tread region and the step region (Figure 5E). The ICDs are no longer in parallel with the Z discs, but are slanted and intersect with Z discs. The ICDs also appear less flat and the amplitude of folding is significantly increased. As a consequence, loss of sarcomere structures in the connecting region between myofibrils and intercalated discs are frequently detected (Figure 5E).

### Identification and functional validation of pathogenic variants in the *SORBS2* gene found in human ARVC patients

Given the cardiac-enriched expression of Sorbs2 protein, its subcellular expression pattern in the ICDs, and depletion of which in the *Sorbs2*^*e8/e8*^ mice leads to ARVC-like phenotypes in human including the prominent RV dilation, cardiac arrhythmia and premature death, we postulated that *SORBS2* is a previously unrecognized gene for ARVC. To test this hypothesis, we performed targeted sequencing as well as Sanger sequencing in a 59 unrelated Han Chinese ARVC patient cohort and WES in 402 health controls, aiming to identify potential pathogenic variants in the coding region of the *SORBS2* gene. Patients with ARVC are significantly younger and have larger right ventricular interior diameter (RVID), right atrial interior diameter (RAID) than health controls (Table 2 in the online-only Data Supplement). Five rare variants were identified in the *SORBS2* gene from five unrelated ARVC patients diagnosed with the task force criteria for ARVC (Figure 6B, Table 2, and Figure 6, Table 3, Table 4 in online-only Data Supplement),^32^ two of them (c.679+1G>T, c.869+1C>G) considered pathogenic according to the American College of Medical Genetics (ACMG) guidelines. By contrast, one likely pathogenic variant (c.3287T>C, p.Val1096Ala) was found in 402 health controls. None of these three variants were reported in human gene mutation database (HGMD) or ClinVar databases. Chi-square test showed potential pathogenic variants in the *SORBS2* gene were significantly enriched in ARVC patients (p=0.0444). Both pathogenic variants are splicing variants predicted to affect splicing donors and/or acceptors, potentially leading to functional disruption of the SORBS2 protein in the patients with these mutations. Among these two ARVC patients, one patient also harbors a nonsense variant in *PKP2* gene (c.2421C>A, p.Tyr807Ter), a known ARVC causative gene (Table 3 in online-only Data Supplement). The other patient did not carry any extra potential pathogenic variants for known ARVC causative genes, except for the c.679+1G>T. The patient carrying this c.679+1G>T splicing variant was a 42 year-old man who had palpitation during exercise. Ventricular tachycardia was detected by electrocardiogram; T wave inversion in the right precordial leads (V_1_, V_2_ and V_3_); RV dilation and fatty filtration were confirmed by MRI (Figure 6B, 6C).

**Table 2.** Summary of *SORBS2* variants identified in 59 ARVC patients

| Protein Change | Nucleotide Change | Polyphen-2 (Score) | SIFT (Score) | 1000 Genome (%) | GnomAD (%) | ExAC (%) | ESP (%) |
| --- | --- | --- | --- | --- | --- | --- | --- |
| Splicing | c.679+1G>T | N/A | N/A | 0.00 | 0.00 | 0.00 | 0.00 |
| Splicing | c.869+1C>G | N/A | N/A | 0.00 | 0.00 | 0.00 | 0.00 |
| p.His745inv | c.2232_2233inv_GTG | N/A | N/A | 0.00 | 0.00 | 0.00 | 0.00 |
| p.Ala1148Val | c.3443G>T | Benign (0.07) | Damaging (0.038) | 0.0016 | 0.00 | 0.0005 | 0.00 |
| p.Met1171Val | c.3511T>C | Damaging (0.993) | Tolerate (0.129) | 0.00 | 0.00 | 0.00 | 0.00 |

PolyPhen-2, Polymorphism Phenotyping v2; SIFT, Sorting Intolerant from Tolerant; GnomAD, Genome Aggregation Database; ExAC, Exome Aggregation Consortium; ESP, Exome Sequencing Project ; N/A, not available.

**Figure 6.**
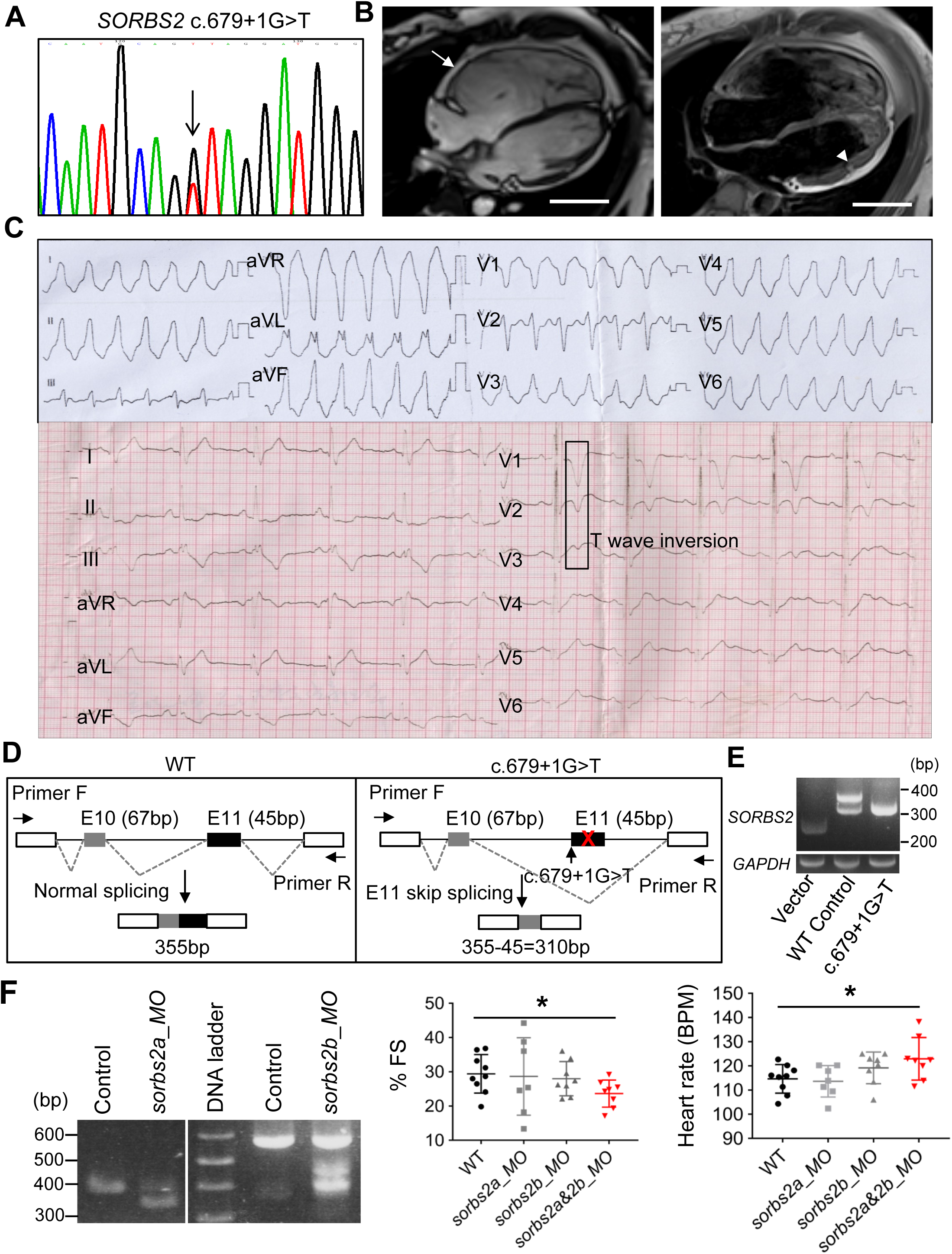
Identification and functional validation of a *SORBS2* splicing variant found from a human patient with ARVC. **A**, Chromagraph illustrates the c. 679+1 G>T mutation identified in *SORBS2* gene from an ARVC patient. **B**, MRI images of the patient harboring the c. 679+1 G>T mutation indicate RV dilation (arrow) and fatty infiltration (arrowhead). Scale bars, 5 cm. **C**, Electrocardiograms from the ARVC patient harboring the c.679+1G>T mutation. Ventricular tachycardia and T wave inversion were noted. **D**, Schematics of the minigene and RT-PCR analysis to test that the splicing mutant c. 679+1G>T leads to exon 11 skip in the HEK 293 cell culture system. **E**, RT-PCR assessment of molecular lesions induced by *sorbs2a* or *sorbs2b_* morpholino (MO) in zebrafish embryos. A predicted 352 bp PCR band was detected in embryos injected with *sorbs2a*_*MO* due to an exon skipping effect. A predicted 447 bp PCR band was detected in in embryos injected with *sorbs2b* MO due to an intron retention effect. **F**, Fractional shortening (FS) and heart rate analysis in *sorbs2a_MO, sorbs2b_MO, sorbs2a&2b_MO* injected morphants and uninjected controls. N=7-9. One-way ANOVA. *: p< 0.05. MO: morpholino.

To validate the pathogenicity of the c.679+1G>T splicing variant, we first conducted mini gene hybridization assay and confirmed that the c.679+1G>T mutation is able to induce exon skipping events (Figure 6D). Loss of exon 11 putatively disrupts the Sorbin homologue domain and results in SORBS2 protein loss-of-function. To further assess the pathogenicity of the c.679+1 G>T point mutation at the organ level, we mimic the genetic lesion in a zebrafish embryo by using the morpholino technology. The targeted sites are exon 6 in *sorbs2a* (ENSDART00000138475.2) and exon 10 in *sorbs2b* (ENSDART00000142605.3), two corresponding exons to exon 11 in human SORBS2, as evidenced by remarkable sequence conservation in the surrounding exons (Figure 7 in the online-only Data Supplement). Injection of *sorbs2a* morpholino caused skipping of exon 6 in *sorbs2a* RNA transcript, while injection of *sorbs2b* morpholino caused intron retention, resulting in a premature stop codon that presumably leads to truncation of the Sorbs2b protein (Figure 6D, 6E). While injection of *sorbs2a* and *sorbs2b* morpholino alone did not cause obvious abnormalities, co-injection of both *sorbs2a* and *sorbs2b* morpholino led to significantly decreased fraction shortening (FS) and increased heart rate (Figure 6F), confirming the pathogenicity of this splicing variant.

## DISCUSSION

### 1. *SORBS2* is a new susceptibility gene for ARVC

Prompted by our genetic studies in zebrafish, we studied cardiac expression and function of *Sorbs2* in mouse. We noted its cardiac enriched expression pattern and the desmosomal/ICD subcellular localization within cardiomyocytes. Homozygous *Sorbs2*^*e8/e8*^ mice manifest characteristic features of other mouse ARVC models, such as those based on *Dsg2, Jup* and *Dsp* genes ^12, 33-35^, including right ventricle (RV) remodeling and dysfunction ^12, 35, 36^, spontaneous sustained and non-sustained ventricular tachycardia ^32^, and premature sudden death at about 3.5 months of age. Importantly, two pathogenic mutations (splicing variants) out of five rare variants in the *SORBS2* gene were identified from a cohort of 59 Han Chinese patients, while only one likely pathogenic variant (missense) was identified in 402 healthy controls. Functional validation studies in the zebrafish model confirmed the pathogenicity of one splicing variant. These data suggest that pathogenic mutations, most likely loss-of-function variants in *SORBS2* gene, are associated with ARVC. Together, these studies in both animal models and human genetics strongly suggested *SORBS2* as an ARVC susceptibility gene, pathogenic mutations in which account for about 3.4% (2 out of 59) of the ARVC patients. Scanning *SORBS2* mutations in larger patient cohorts are needed to accurately determine its prevalence.

Different from other AVRC genes that were mostly discovered through human genetic studies, *SORBS2* is the first ARVC gene that was initially identified from expression and phenotypic studies in animal models. We acknowledged limitations associated with our study. Fatty infiltration, a pathogenic hallmark for human ARVC, was not prominent in our *Sorbs2*^*e8/e8*^ mouse model. Similarly, it does not present in most of the mouse ARVC models for other well-established ARVC genes such as *JUP, DSG2* and *PKP*, neither^12, 33^. The only mouse ARVC model with fibro-fatty infiltration phenotype is the Dsp-deficient mouse ^12, 13^. Species difference has been postulated to explain the discrepency, becuase a mouse heart does not show evidence of epicardial fat as typically seen in a human heart. Moreover, the present human genetic studies were conducted in sporadic ARVC patients. Pedigree studies are needed to further validate *SORBS2* as an ARVC causative gene.

### 2. *Sorbs2*^*e8/e8*^ is a mouse model for *SORBS2* ARVC

The *Sorbs2*^*e8/e8*^ mouse provides a useful animal model to decipher the mechanisms of SORBS2 ARVC. In ARVC patients, sudden cardiac death (SCD) and ventricular arrhythmia are often seen at the early phase when there is no obvious heart structural remodeling ^37^. Similar progressive pathogenesis was observed in the *Sorbs2*^*e8/e8*^ mice. Spontaneous and induced ventricular arrhythmias are noted as early as 2 months of age in the *Sorbs2*^*e8/e8*^ mice (data not shown). RV remodeling occurs later, followed by LV hypertrophy. Electrophysiology experiments identified multiple levels of abnormalities. First, the ECG abnormalities with prolonged and bifid P waves and RBBB consistent with atrial and ventricular conduction abnormalities. Second, spontaneous atrial and ventricular arrhythmias were recorded in anesthesized *Sorbs2*^*e8/e8*^ mice. Third, microelectrode studies showed ventricular depolarized resting potentials, reduced action potential upstroke dV/dts, reduced action potential amplitudes, and prolonged ADP50s, ADP90s and VERPs, indicating potential derangements in a multitude of ion channels including those that encode IK_1_, I_Na_, I_to_, and I_K_. Fourth, Sorbs2 deficient hearts are susceptible to induction of VT, which may involve reentry and triggered activity in mechanisms. These findings indicate that the loss of Sorbs2 leads to loss of structural integrity and is accompanied by a wide range of electrophysiological abnormalities that promote development of arrhythmias and premature death.

Compared to other well-established ARVC susceptibility genes that encode desmosome proteins, *SORBS2* encodes an ICD protein that is predominantly localized to the adhesion junction but imparts its effects in gap junctional proteins. Nevereless, similar ICD structural disruption was noted in the *Sorbs2*^*e8/e8*^ mice, as often seen in ARVC mouse models and human patients, including breakdown of the ICD structure, a widened desmosome gap, and mislocated desmosome ^35, 38^. We also noted the loss of myofibrils in the proximity of the ICD region, as has been reported in Boxer dogs with ARVC ^39^. Sorbs2 depletion did affect the expression pattern of other ICD proteins, such as N-cadherin, also known as cadherin-2 (Cdh2), which is encoded by a recently identified ARVC gene ^9^. How Sorbs2 protein interacts with other ICD proteins, and whether Sorbs2 deficiency regulates known ARVC pathways such as Wnt, YAP and/or mechanical force transmission warrants future investigation^12, 40, 41^. Intriguingly, Sorbs2 deficiency is associated with dramatical downregulation of Cx43 expression in mouse. Given the fact that marked reduction of Cx43 have been found in many ARVC patients ^28 42, 43^, it would be interesting to investigate the underlying mechanism its contribution to arrhythmic phenotypes in ARVC.

In summary, our integrated studies from zebrafish, mouse and human genetics suggested *SORBS2* as a new susceptibility gene for ARVC, which can be modeled by a *Sorbs2* KO mouse.

## Acknowledgements

We thank Dr. Prasanna Mishra in the Mayo Nuclear Magnetic Resonance lab for guidance on MRI in mouse, Ronald May for performing echocardiography in mice, Mayo Clinic Electron Microscopy Core Facility for transmission electron microscopy in mice, and Mayo Histology Core Facility for performing masson’s trichrome staining.

## Sources of Funding

This work was supported by grants from NIH R01-HL107304, R01-HL081753, R01-GM63904 and Mayo Foundation to XX; The Innovative Basic Science Award (1-16-IBS-195) from American Diabetes Association and the Prospective Research Award from the Mayo Clinic Department of Cardiovascular Diseases to TL; NIH R01-HL074180 and Mayo Foundation to HL.

## Disclosures

None.

