## Supplemental Figures, Tables and Methods for "*SORBS2* is a susceptibility gene to arrhythmogenic right ventricular cardiomyopathy"

### Supplementary information

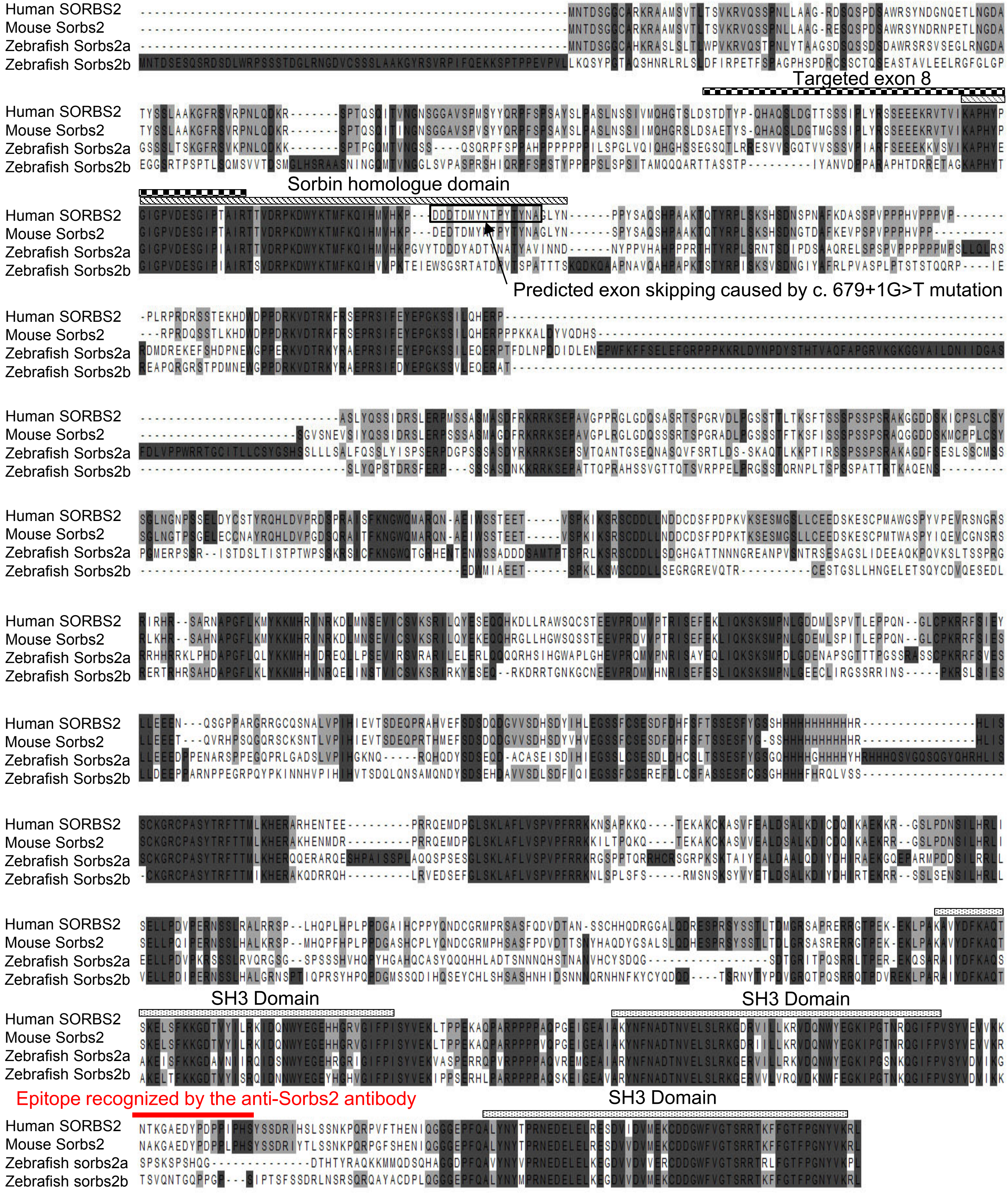


**Supplementary Figure 1.** Alignment of Sorbs2 protein sequences from human, mouse and zebrafish. Sequences are more conserved in the N-terminal sorbin homologue domain and the C-terminal SH3 domain among 3 different species. The deletion in *Sorbs2* $^{e8/e8}$ mice removed N terminal sequences containing part of the sorbin homologue domain. The 15 amino acids that are encoded by exon 11 and disrupted by 679+1 G>T mutation are enclosed by a black rectangle frame. Epitope region recognized by the anti-Sorbs2 antibody used for Western blot and immunostaining in mouse tissues is indicated by a red solid line.

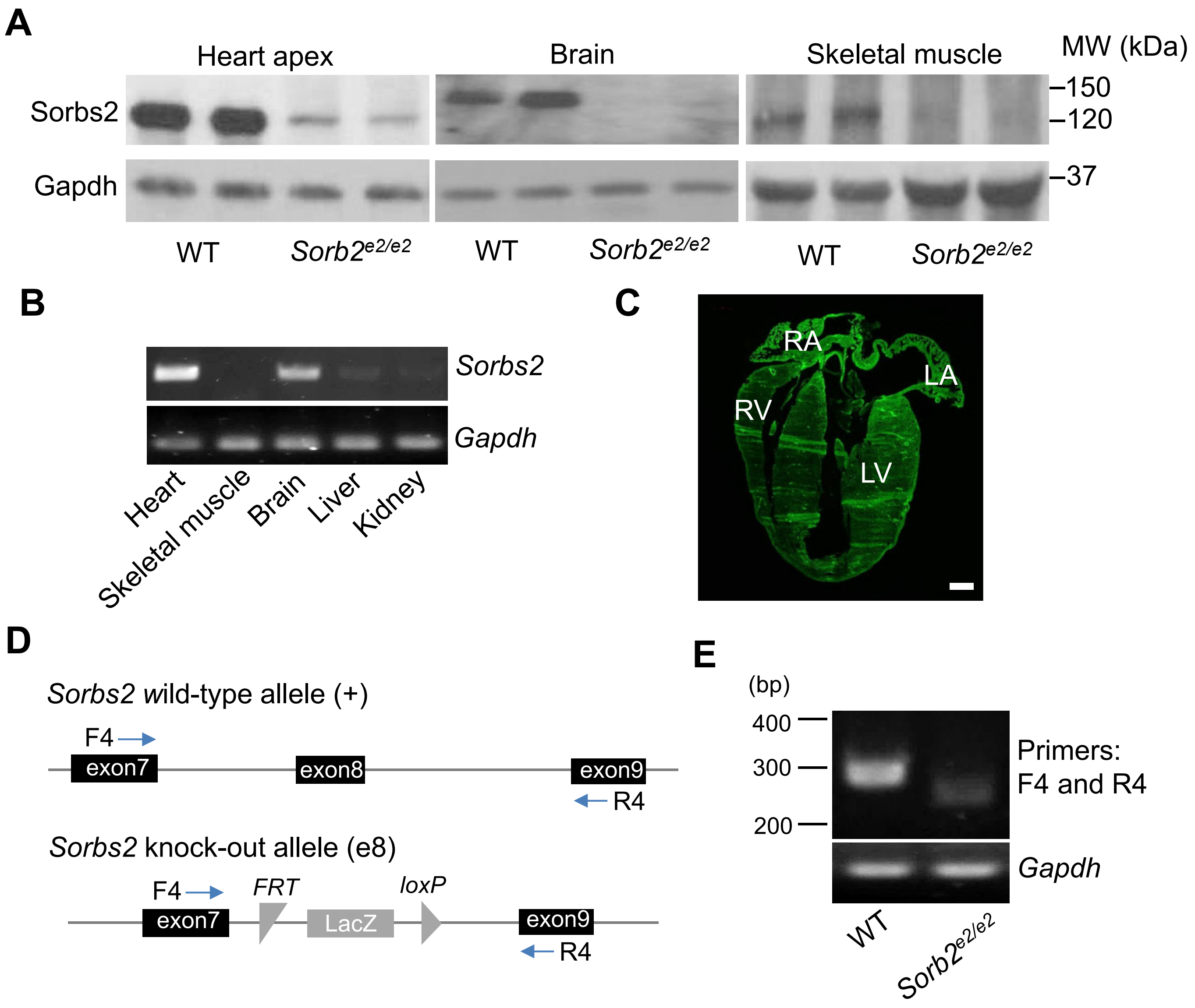


**Supplementary Figure 2.** Targeted depletion of the exon 8 in the *Sorbs2*$^{e8/e8}$ mice**. A,** Western blotting analysis of Sorbs2 protein expression in the heart, brain and skeletal muscle of *Sorbs2*$^{e8/e8}$ and WT control mice. **B,** RT-PCR analysis of the *Sorbs2* transcript expression in different mice tissues. **C,** Shown are immunostaining of mice heart sections using an anti-Sorbs2 antibody. Sorbs2 is ubiquitously expressed in 4 cardiac chambers and septum. Scale bar: 50 µm. **D,** Schematic design of primers for RT-PCR to analyze the targeted deletion of exon 8 in the *Sorbs2*$^{e8/e8}$ mice. **E**, Shown are representative DNA gel images of RT-PCR to confirm the replacement of exon 8 with LacZ that leads to frameshift of Sorbs2 transcript in the *Sorbs2* $^{e8/e8}$ mice.

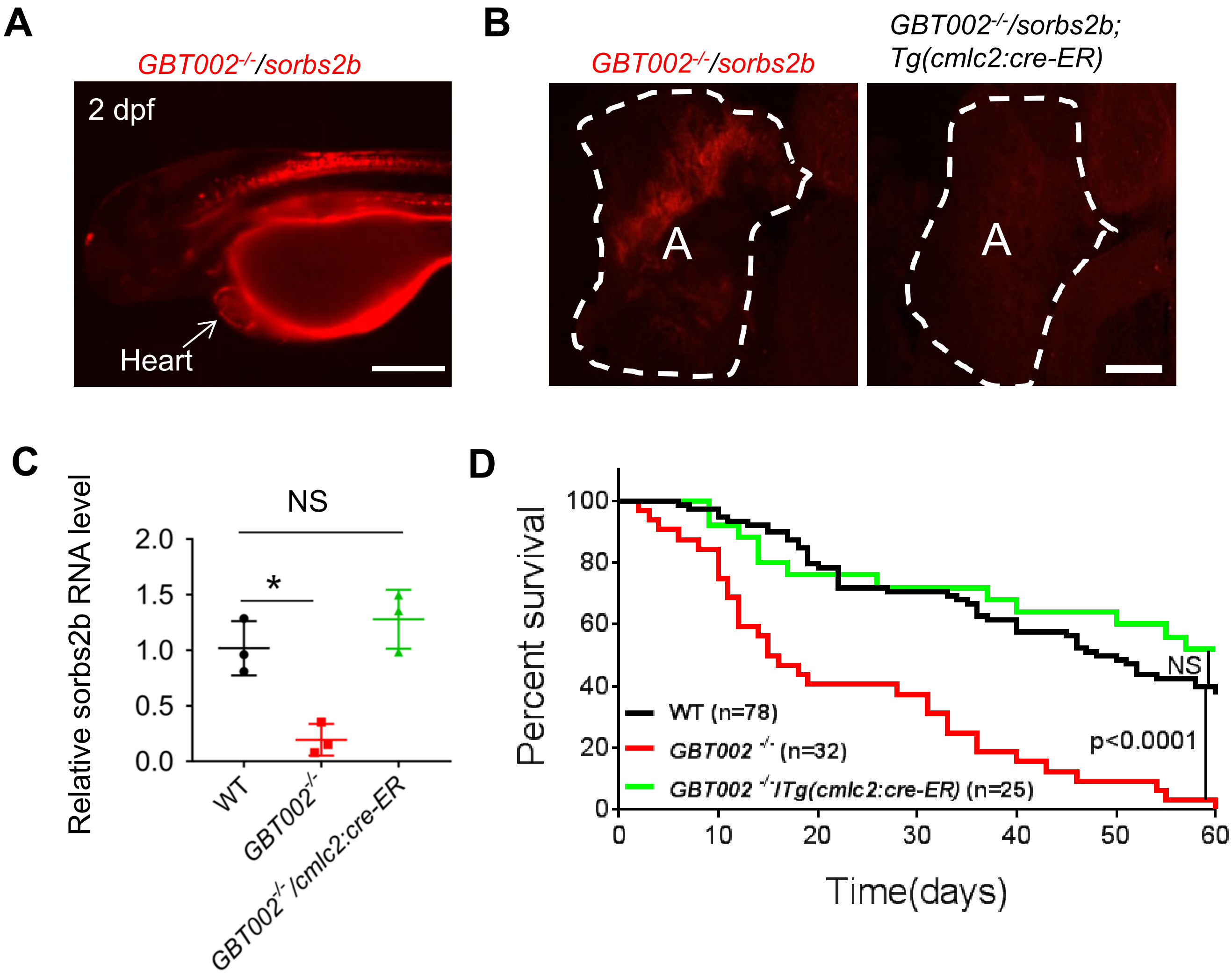


**Supplementary Figure 3.** Myocardial expression of *sorbs2b* conveys the modifying effects of *GBT002*$^{-/-}$ on doxorubicin-induced cardiomyopathy (DIC) in zebrafish. **A,** Heart enriched expression of red fluorescent protein (RFP) reporter in the *GBT002/sorbs2b* mutant at 2 days post-fertilization (dpf). Scale bar, 200 µm. **B,** RFP expression in the *GBT002*$^{-/-}$ mutant heart, representing an RP2 insertion in the sorbs2b locus, was effectively excised in the *GBT002*$^{-/-}$*;Tg(cmlc2:cre-ER)* double fish after hydroxytamoxifen treatment. Scale bar: 200 µm. **C**, Quantitative RT-PCR results show that normal *sorbs2b* RNA is dramatically reduced in *GBT002*$^{-/-}$ fish compared to WT, but was largely restored in the *GBT002*$^{-/-}$*; Tg(cmlc2:cre-ER)* double fish heart. N=3. * : $p<0.05$. NS, not significant. One-way ANOVA. **D**, The survival rate is significantly lower in *GBT002*$^{-/-}$ mutant compared to WT fish after doxorubicin injection (20 µg/g) , which was largely rescued in the *GBT002*$^{-/-}$*; Tg(cmlc2:cre-ER)* double fish. N=25-78. NS, not significant. Log rank test.

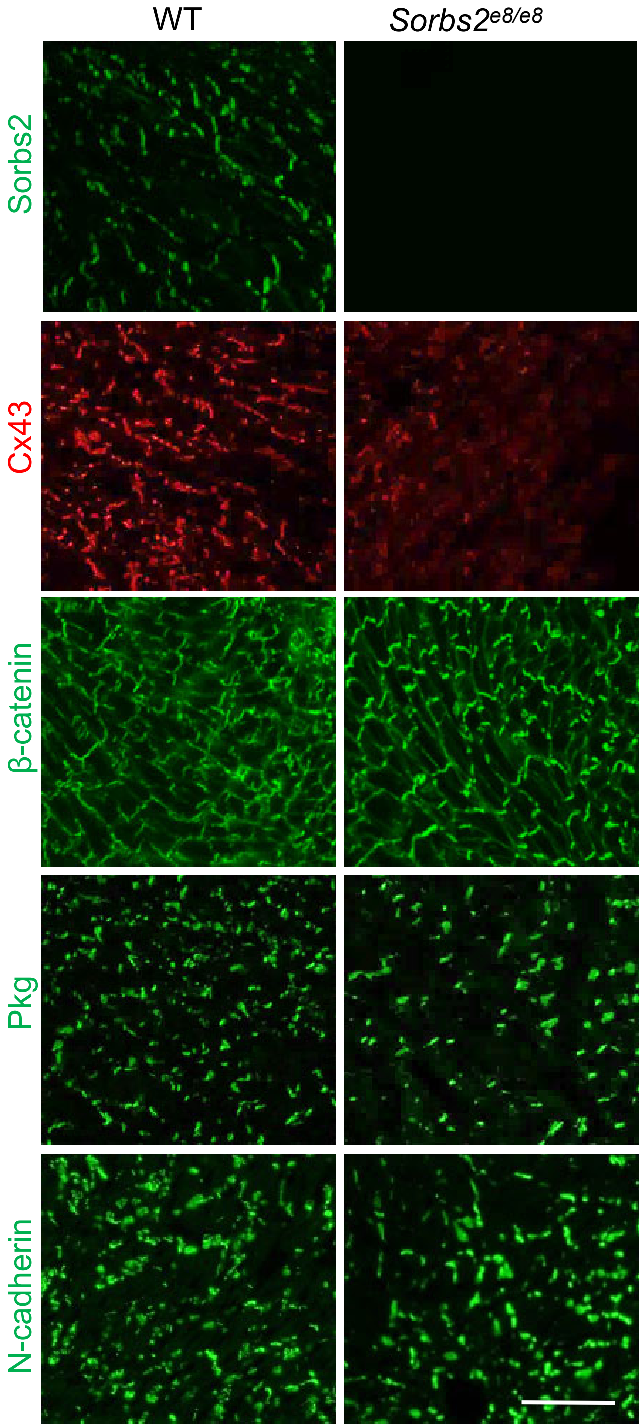


**Supplementary Figure 4.** Representative Immunostaining images of ICD proteins in the *Sorbs2*$^{e8/e8}$ and WT control mice mouse hearts. Expression levels of Sorbs2 and Cx43, but not other ICD proteins are reduced in the *Sorbs2*$^{e8/e8}$ mice.. Scale bar: 100 μm.

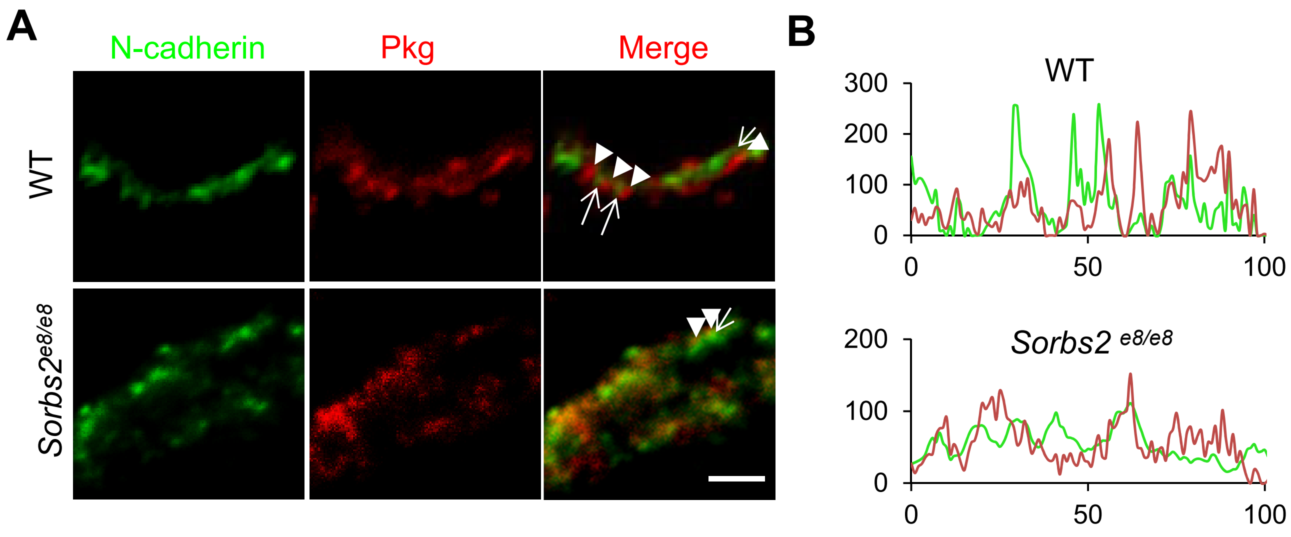


**Supplementary Figure 5.** Altered expression patterns of N-cadherin in the *Sorbs2* $^{e8/e8}$ mice. **A**, Shown are images of mouse hearts after immunostaining using anti-N-cadherin antibodies (green, indicated by arrows) and anti-Pkg (red, indicated by arrow heads). The alternating expression pattern of Pkg and N-cadherin protein expression in WT mice is disturbed in *Sorbs2* $^{e8/e8}$ mice along ICD. Scale bar: 2 μm. LV: left ventricle, RV: right ventricle. **B,** Pseudo line analysis of images in **A**.

**A**

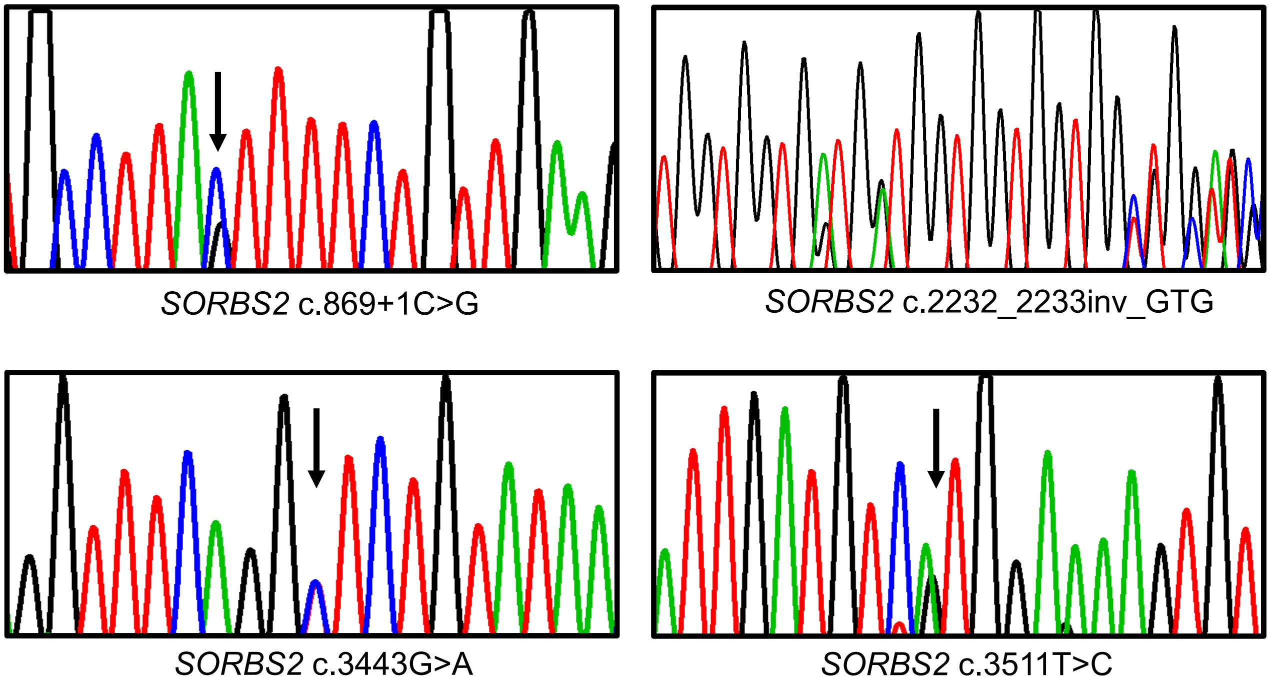


**B**

| No. | Gene | Official Full Name | Chr. | CDS n | Amplicons n | Target bp | Coverage % |
|---|---|---|---|---|---|---|---|
| **1** | *DES* | Desmin | 2 | 9 | 17 | 1512 | 100 |
| **2** | *DSC2* | Desmocollin 2 | 18 | 17 | 31 | 2606 | 89.0 |
| **3** | *DSG2* | Desmoglein 2 | 18 | 15 | 36 | 3420 | 97.1 |
| **4** | *DSP* | Desmoplakin | 6 | 25 | 74 | 9364 | 99.7 |
| **5** | *JUP* | Junction Plakoglobin | 17 | 13 | 22 | 2381 | 100 |
| **6** | *LMNA* | Lamin A/C | 1 | 13 | 23 | 2249 | 100 |
| **7** | *PKP2* | Plakophilin 2 | 12 | 14 | 28 | 2640 | 94.3 |
| **8** | *PLN* | Phospholamban | 6 | 1 | 10 | 1139 | 75.8 |
| **9** | *RYR2* | Ryanodine Receptor 2 | 1 | 105 | 165 | 14830 | 99.5 |
| **10** | *SCN5A* | Sodium Voltage-Gated Channel Alpha Subunit 5 | 3 | 30 | 62 | 6807 | 100 |
| **11** | *TGFB3* | Transforming Growth Factor Beta 3 | 14 | 7 | 11 | 1316 | 100 |
| **12** | *TMEM43* | Transmembrane Protein 43 | 3 | 12 | 16 | 1335 | 100 |
| **13** | *TTN* | Titin | 2 | 315 | 947 | 109511 | 99.1 |

**Supplementary Figure 6.** Identifications of *SORBS2* variants in 59 ARVC patients. **A,** Chromatographs of other four variants in *SORBS2* identified in four different ARVC patients. **B,** List of the ARVC gene panel that consists of 13 genes. Sequence variants were searched by targeted sequencing.

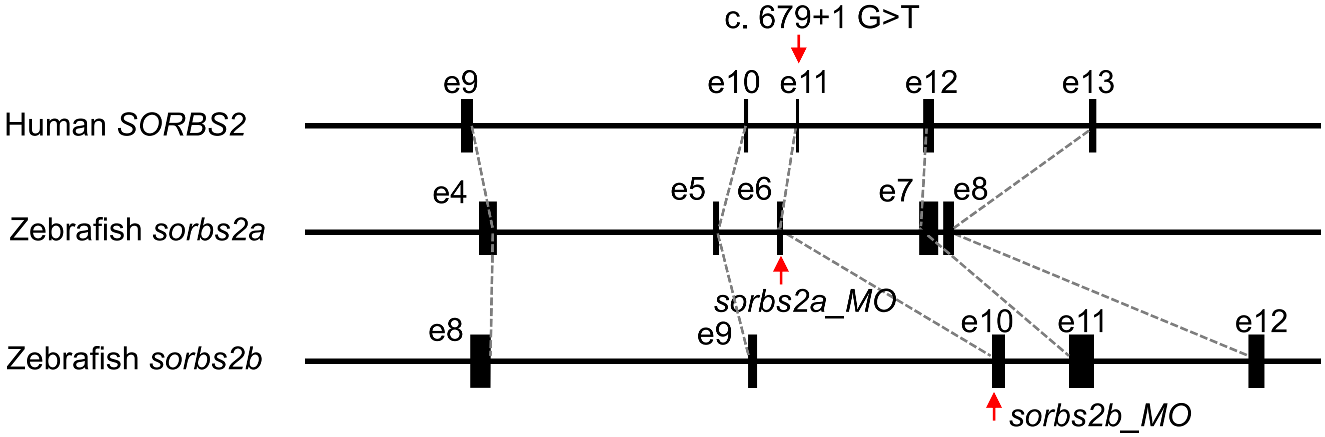


**Supplementary Figure 7.** Schematic illustration of corresponding exons among human *SORBS*2 and its zebrafish orthologs *sorbs2a* and *sorbs2b* genes. The c.679+1G>T mutation is located in the exon 11 of *SORBS2* in human, corresponding to the exon 6 in *sorbs2a* and the exon 10 in *sorbs2b* gene in zebrafish.

| | WT siblings | *Sorbs2*$^{e8/-}$ | P value |
|---|---|---|---|
| Mice number (n) | 15 | 14 | |
| HR (bpm) | 356±63 | 350±74 | 0.829393 |
| IVSd (mm) | 0.72±0.07 | 0.70±0.05 | 0.363821 |
| LVIDd (mm) | 3.90±0.29 | 4.03±0.42 | 0.333019 |
| LVPWd (mm) | 0.79±0.27 | 0.70±0.07 | 0.22288 |
| IVSs (mm) | 1.10±0.11 | 1.08±0.09 | 0.518077 |
| LVIDs (mm) | 2.79±0.29 | 2.92±0.44 | 0.348279 |
| LVPWs (mm) | 1.06±0.13 | 1.05±0.12 | 0.882353 |
| LVEF (%,Cube) | 63.07±5.64 | 63.50±5.82 | 0.843029 |
| LVEF (%, Teich) | 61.73±5.58 | 61.86±5.82 | 0.954625 |
| LVFS (%) | 28.4±3.69 | 28.71±3.84 | 0.82698 |
| LVd Mass (g) | 0.68±0.01 | 0.68±0.02 | 0.742906 |
| LVs Mass (g) | 0.68±0.02 | 0.69±0.02 | 0.807624 |

**Supplementary Table 1.** Echocardiography indices in the *sorbs2*$^{e8/-}$ heterozygous mice compared to WT controls at 4 weeks post-doxorubicin injection (20 mg/kg). HR, heart rate; bpm, beats per minute; IVSd, Interventricular septum thickness at end–diastole; LVIDd, left ventricular internal dimension at end-diastole; LVPWd, left ventricular internal dimension at end-diastole; IVSs, Interventricular septum thickness at end–systole; LVIDs, Left ventricular internal dimension at end-systole; LVPWs, Left ventricular posterior wall thickness at end–diastole; LVEF, left ventricular ejection fraction; LVFS, left ventricular fractional shortening; LVd, left ventricular at end-diastole; LVs, left ventricular at end-systole. Two-paired student's *t*-test.

| | Patients with ARCV | Healthy Control | p-value |
|---|---|---|---|
| | (n=59) | (n=402) | |
| Age, y | 47.7 ± 12.4 | 51.8 ± 10.8 | <0.0001 |
| Male, n (%) | 64 (62.7) | 330 (66.0) | 0.5293 |
| Hypertension, n (%) | 32 (31.4) | 286 (57.2) | <0.0001 |
| Diabetes, n (%) | 12 (11.8) | 93 (18.6) | 0.0971 |
| Alcohol, n (%) | 21 (20.6) | 113 (22.6) | 0.6562 |
| LAID, mm | 47.4 ± 11.1 | 33.5 ± 5.6 | <0.0001 |
| LVIDd, mm | 55.7 ± 8.8 | 47.7 ± 4.7 | <0.0001 |
| RAID, mm | 57.2 ± 9.2 | 38.7 ± 5.9 | <0.0001 |
| RVIDd, mm | 53.4 ± 10.1 | 29.1 ± 4.3 | <0.0001 |
| Maximum LV wall thickness, mm | 9.72 ± 1.71 | 9.2 ± 0.7 | 0.3124 |
| Left ventricular eject fraction, % | 48.5 ± 13.2 | 65.2 ± 8.6 | <0.0001 |
| Implantable cardioverter-defbrillators, n (%) | 21 (20.6) | 0 (0.0) | <0.0001 |

**Supplementary Table 2.** Baselines of 59 ARVC patients and 500 health control. LAIDd, left atrial internal dimension at -diastole; LVIDd, left ventricular internal dimension at -diastole; RAIDd, right atrial internal dimension at diastole; RVIDd, right ventricular internal dimension at diastole.

**Supplementary Table 3.** Clinical Characteristics and Genotypes of the Patients Carrying Potential Pathogenic Variants in the *SORBS2* and other 14 known ARVC genes.

| Gene | Genotype | Age, y | LAIDd, mm | LVIDd, mm | RAIDd, mm | RVIDd, mm | LVEF, % | Mutations for other 14 known ARVC genes |
|---|---|---|---|---|---|---|---|---|
| *SORBS2* | c.679+1G>T | 43 | 32 | 50 | 33 | 45 | 68 | None |
| *SORBS2* | c.869+1C>G | 27 | 29 | 42 | 47 | 49 | 66 | *PKP2*: c.2421C>A |
| SORBS2 | c.2232_2233inv_GTG | 47 | 33 | 36 | 48 | 63 | 63 | None |
| *SORBS2* | c.3443G>T | 86 | 40 | 39 | 65 | 61 | 63 | None |
| *SORBS2* | c.3511T>C | 30 | 37 | 48 | 55 | 43 | 59 | *DSP*: c.861T>G, c.8605A>G<br>*DSG2*: c.2377A>G |

RefSeq: *SORBS2*: NM_001270771.1; LAIDd, left atrial internal dimension at diastole; LVIDd, left ventricular internal dimension at diastole; RAIDd, right atrial internal dimension at diastole; RVIDd, right ventricular internal dimension at diastole; LVEF, left ventricular ejection fraction. ARVC, arrythmogenic right ventricle cardiomyopathy.

**Supplementary Table 4.** Diagnosis of the Patients Carrying Potential Pathogenic Variants in *SORBS2* gene according to Task Force Criteria (Marcus et al. Circulation 2010)

| Genotype | Global and/or Regional Dysfunction and Structural Alterations | Tissue Characterization of Wall | Repolarization Abnormalities | Depolarization/ Conduction Abnormalities | Arrhythmias | Family History |
|---|---|---|---|---|---|---|
| c.679+1G>T | Major | NA | Major | Minor | Major | NA |
| c.869+1C>G | Major | NA | Major | Minor | Major | NA |
| c.2232_2233inv_GTG | Major | NA | Minor | - | Minor | NA |
| c.3443G>T | Major | NA | Major | Minor | Minor | NA |
| c.3511T>C | Major | NA | Minor | Minor | Minor | NA |

NA, data not available

**upplementary Table 5.** Sanger sequencing primers of coding regions of the *SORBS2* gene

| e | Forward | Reverse |
|---|---|---|
| BS2-E4 | 5'- AAGAGTGGGGCACAAATACAAAG-3’ | 5'- AAACAAGCACAAAACGTCTCCAT-3’ |
| BS2-E5 | 5'- AAACGTGAATAGTTATTAACACTCGC-3’ | 5'- GGTGCATTCCATGATTACTAATTCT-3’ |
| BS2-E6E7 | 5'- GAAAGAGGCTGTAAGCAGTGGTC-3’ | 5'- TTGGCATGTTTGACATACCCA-3’ |
| BS2-E8 | 5'- AGACACAGGTGGAGACATGGTCT-3’ | 5'- TACTCTCATGTGATTGGAAGTCTTTTA[illegible] |
| BS2-E9 | 5'- AGGAATGCACAAAGCCGTG-3’ | 5'- CTCTGTGTTACAGTTGGCATAAATATA[illegible] |
| BS2-E10 | 5'- AACCGATGCAAAAGACGAATG-3’ | 5'- GCAAAGAGTCAGAACATTTTATCACA-[illegible] |
| BS2-E11 | 5'- CCTTTTGTTATGCACGCTCTGA-3’ | 5'- TACGTTGCTTTTCTAACAAATTTCTG-3[illegible] |
| BS2-E12 | 5'- TGGCAGGTGCACTGATTCTTC-3’ | 5'- CAAAGATTTACTGTAATCAGGGCTTA-[illegible] |
| BS2-E13 | 5'- AAATTGCTAATGGGTCTGAAATTC-3’ | 5'- CTCATGTGACTTTTGGAGCCAG-3’ |
| BS2-E14 | 5'- TAAGGGCCATTTAACAACATTGA-3’ | 5'- CCAAGTAGAAATCAGGTCGCAT-3’ |
| BS2-E15 | 5'- ATTCCAGACACTGTAGGTGAGAGC-3’ | 5'- GGTTTAGGAGTTACGCCATTGTC-3’ |
| BS2-E16-Part1 | 5'- GGGAGTGATGAGATGTTGGAATG-3’ | 5'- ACATTGAAGTCACCAGCGATGA-3’ |
| BS2-E16-Part2 | 5'- GTAAATCGAGTGTAGGAGGCCG-3’ | 5'- GAAGCTCCATGTGTGAACAATCA-3’ |
| BS2-E17 | 5'- GAGTTCGTTACTGGTTTGGGATA-3’ | 5'- TGTGTTCGATAACGGTGTCTCTC-3’ |
| BS2-E18 | 5'- GATCCTACTGCAATGCTTATCTCTTA-3’ | 5'- GTTTGATGATTCCCTGAGGTTTC-3’ |
| BS2-E19E20 | 5'- ACCAGCAGGGAACCGTGAC-3’ | 5'- CAAGGATCATTCTTATCTAATCGGAC-[illegible] |
| BS2-E21 | 5'- TTGTCAACCTAATAACCCGGTG-3’ | 5'- TCTGATGGCCTGAATGGACAT-3’ |
| BS2-E22 | 5'- CGTGCAAAGCCATACATCTTACA-3’ | 5'- ACATTTGACCTTGCTTATTTTGCT-3’ |
| BS2-E23 | 5'- GCCATCACAGCAAACCACG-3’ | 5'- ACAATTTTCATGGATATTCCTCGT-3’ |
| BS2-E24 | 5'- TTGGAAACCGTAAGGCATGA-3’ | 5'- CACTGCTTATTTTGTCTGCCTATG-3’ |

#### Expanded Online Methods

##### Animal Experiments

All procedures were carried out in accordance with Mayo Clinic guidelines for animal use and care and conformed to the Guide for the Care and Use of Laboratory Animals published by the US National Institutes of Health. *Sorbs2$^{e8/e8}$* mice, also named *B6N(Cg)-Sorbs2 $^{tm1.1(KOMP)Mbp/J}$* were purchased from the Jackson Laboratory (Catalog #022780) with genotype characterization using PCR primers: 5'-ATTAGGACGGTAAGCCACGC-3' and 5'-AGGCTTCACCTTGCTTTGATAG-3' for detecting wild type (WT) allele (predicted size of 292 bp), and using primers: 5'-TTTGGCTTCAACCACAACAC-3' and 5'-CGGTCGCTACCATTACCAGT-3' for detecting mutant allele (predicted size of 500 bp). Zebrafish (Danio rerio) WIK line was maintained under a 14 hours light/10 hours dark cycle at 28.5°C. WT zebrafish were bred to give embryos for morpholino injection and subsequent fractional shortening and heart rate analysis. Animal study protocols approved by the Mayo Clinic Institutional Animal Care and Use Committee for mouse is IACUC A00002911, and for zebrafish is A3531.

##### Tissue collection

For tissue harvest, mice were sacrificed by administration of 250 mg/kg pentobarbital, according to the guidelines of the Mayo Clinic Institutional Animal Care and Use Committee. Heart tissues from different cardiac chambers were frozen in liquid nitrogen and stored at -80°C and were used for total RNA transcript and western-blot experiments. Some tissue were fixed in formalin and embedded in paraffin for histomorphometry. At least 3 heart samples were collected for each individual experiment.

##### Western-blot

Heart tissues were homogenized in RIPA buffer with 0.5 mm steel beads at speed 8 for 4 minutes followed by speed 10 for 1 minute using a Bullet Blender (Next Advance). Lysate samples were denatured at 95°C for 10 min and loaded onto a sodium dodecyl sulfate–polyacrylamide gel for electrophoresis (SDS-PAGE) gel and Western blotting. Anti-Sorbs2 (Sigma, Cat#SAB4200183), Anti-Cx43 (Cell Signaling, Cat#3512), anti-β-Catenin (Thermofisher Scientific, Cat#13-8400), anti-Pkg (Progen, Cat#651167), anti-N-Cadherin (Sigma, Cat#C1821), and anti-Gapdh (Santa Cruz Biotechnology, Cat#sc-25778) antibodies were used.

**Quantitative RT-PCR**

Total RNA was extracted using the RNeasy extraction kit (Qiagen) according to manufacturer's instructions. The extracted RNA was quantified and 1 μg RNA was reverse transcribed to synthesize cDNA using the Super Script III First Strand Reverse Transcriptase Kit and random hexamer primers (Invitrogen). After initial denaturation with LightCycler 480 Probes Master mix (Roche), 50 ng of the cDNA underwent 50 rounds of amplification (LightCycler 480, Roche) with the following primers: 5'-GTGCGGTGTCCAACACAGAT-3' and 5'-TCCAATCCTGTCAATCCTACCC-3' for mouse *Nppa* gene; 5'-GAGGTCACTCCTATCCTCTGG-3' and 5'-GCCATTTCCTCCGACTTTTCTC-3' for mouse *Nppb* gene; 5'-ACTGTCAACACTAAGAGGGTCA-3' and 5'-TTGGATGATTTGATCTTCCAGGG-3' for mouse *β-Mhc* gene; 5'-GCCCAGTACCTCCGAAAGTC-3' and 5'-GCCTTAACATACTCCTCCTTGTC-3' for mouse *α-Mhc* gene; 5'-CTGAATGGTATCGCACCGGAA-3' and 5'-GGTCCACAAGCACTCCACAG-3' for mouse *Cx40/Gja5* gene, 5'-ACAGCGGTTGAGTCAGCTTG-3', 5'-GAGAGATGGGGAAGGACTTGT-3' for mouse *Cx43/Gja1* gene, 5'-CAGAGCCAACCAAAACCTAAGC-3', 5'-CTGCACACATAAAATGGGTGGA-3' for mouse *Cx45/Gjc1* gene; and 5'-CTATCAGAGGCCATTCTCCC-3' and 5'-GAACATTGTCTTGTACCAGTCC-3' for mouse *Sorbs2* gene. Data were analyzed and relative expression determined using the comparative cycle threshold (Ct) method (2−ΔΔct), and expression was normalized using *Gapdh (*5'-AGGTCGGTGTGAACGGATTTG-5' and 5'-TGTAGACCATGTAGTTGAGGTCA-3*')* as the housekeeping internal control.

**Immunohistochemistry**

The heart samples harvested from mice were embedded in tissue freezing medium and stored in -80°C , followed by section at 10 μm using a cryostat (Leica CM3050 S). The slides were subjected to immunostaining using a previously described protocol [1]. The following antibodies were used: anti-Sorbs2 (Sigma, Cat#SAB4200183) at 1:200, anti-Cx43 (Cell signaling, Cat#3512) at 1:200, anti-N-Cadherin (Sigma, Cat#C1821), anti-β-Catenin (Thermofisher Scientific, Cat#13-8400) at 1:200, and anti-Pkg (Progen, Cat#651167) at 1:200. Zeiss LSM 780 confocal microscope was used to acquire the image. The tile scan function was used to acquire images covering a complete mouse heart. For each group, 5–10 slides were imaged and analyzed. The distribution of fluorescence density was analyzed by pseudo-line scan using the Zen software (Zeiss). The weighted co-localization of Sorbs2 with other ICD protein was determined using co-localization function in the Zen software.

**Magnetic resonance imaging (MRI)**

Mice were anesthetized by inhalational isoflurane (1.5% -2.5% in oxygen/air) via a nose cone during the imaging procedure. Respiration was continuously monitored during the scan and heart rate was kept at 20-60 breaths per minute by adjusting the concentration of isoflurane. The animal might be kept under anesthesia for more than 2 hours. The body temperature of the animals was maintained by a stream of air conditioned by thermocouple-based system at 32°C. MRI was performed in a Bruker Avance 300 MHz (7 Tesla) vertical bore nuclear magnetic resonance spectrometer equipped with "miniimaging" accessories. The images were acquired with a field of view of 3.4 $cm^2$, slice thickness of 1 mm and an in-plane resolution of 135 micrometers. Images were acquired from both long-axial and short axial view, and analyzed off-line using custom-designed code in Matlab [2], [3]. Right and left ventricular endocardial and epicardia contours were drawn on each of the 10 short-axial or long axial slices. The wall thickness of left ventricle was calculated directly from the contour data. The volume of left ventricle was calculated based on Single-plane Ellipsoid model (SP) using the following equation [4]:

$$volume\ of\ left\ ventricle = \frac{8 \times A \times A}{3\pi \times L}$$

While A was the area of left ventricle in long-axis view and L was the length of left ventricle along long-axis view. Right ventricular volumes were calculated using a formula derived from ellipsoidal shell model (difference-of-ellipsoids model): V = 2/3Pd . The plain of area P was defined by two perpendicular axes of the ellipsoid, and distance d was its third axis perpendicular to area P [4].

**Electrocardiogram (ECG)**

Anesthesia with isoflurane (0.5% to 1.0% v/v) was administered via a nose cone. Mice were placed on an ECG-heater board with 4 paws on individual electrode. The ECG-heater board maintained the body temperature at 37°C and monitored the heart rates. The ECG signal was amplified through an Amplifier (Axon CNS digital 1440 A) and recorded using Chart 5 software. For each mouse, 2 minutes of ECG signal were recorded.

**Electrophysiology**

*Ex vivo* cardiac electrophysiology study was performed in Langendorff-perfused mouse hearts as we have described previously [5]. Briefly, the excised heart was rapidly mounted on the cannulus of a modified Langendorff apparatus and perfused at 3.0 mL/min using a peristaltic pump (P720, Instech Laboratories) with Tyrode's solution (in mM): NaCl 119, KCl 4.8, $KH_2PO_4$

1.2, $MgSO_4$ 1.2, CaCl2 1.0, $NaHCO_3$ 24.9, glucose 10.0, pyruvate 5.0, heparin 1200 U/L, pH=7.4), equilibrated with 95% O2-5% CO2 at 37 °C. A bipolar electrode was placed on the right ventricular epicardium for cardiac pacing. Programmed electrical stimulation was performed using a cardiac stimulator (Bloom-Fisher Medical Technologies) with 2-ms current pulses at a two-time diastolic threshold (100-200 µA). The cycle lengths of stimuli were 10 to 20 ms shorter than those of intrinsic heart beat coupling intervals.

Action potential (AP) recordings of isolated mouse RV were performed using standard microelectrode techniques as previously described [6]. The right ventricle was rapidly excised, placed with the endocardial surface up in a temperature-regulated 5-ml chamber, and continuously superfused with oxygenated Tyrode's solution at 4 ml/min at 37 °C. The tissue was impaled with a micropipette with a tip resistance of 10-25 MΩ when filled with 3M KCl, and the output signals were sampled at 100 kHz without filter by a high-input impedance preamplifier (Duo 773, World Precision Instruments, Sarasota, FL). The cardiac preparations were paced using a stimulator (Bloom Associates, Ltd., Reading, PA) with a stimulus output at twice threshold. The action potentials were recorded at a paced cycle length (CL) of 200 ms and the resting potential, APD50 and APD90 were measured off-line. VERP was measured by a train of stimuli (10~14 S1) at a fixed cycle length of 200 ms, coupled with an extra-stimulus (S2) that was 10-ms shorter than the train and repeated at 10-ms decrements until refractoriness was reached. Burst pacing was performed by trains of 20 pulses with coupling intervals 10 to 20 ms longer than VERP. VTs were induced by programmed electrical stimulation and burst pacing. Sustained VT was defined as wide QRS complex tachycardia with a rate greater than 600 beats/min lasting longer than 30 sec [7].

**Transmission Electron Microscopy**

Tissues from mouse hearts were bathed in relaxation buffer (75 mM KCl, 10 mM imidazole pH 7.2, 2 mM $MgCl_2$, 2 mM EGTA, 1 mM $NaN_3$, 4 mM creatine phosphate) for 1 h, fixed in Trump's fixative (Electron Microscopy Sciences) overnight at 4°C, and then processed at the Mayo Clinic's Electron Microscopy Core Facility. Images were obtained using a Philips CM10 transmission electron microscope (Philips, Amsterdam, The Netherlands).

**Hematoxylin-eosin (H&E) and Masson's trichrome staining**

Heart tissues were harvested from mice at 4 months of age after being euthanized by by administration of 250 mg/kg pentobarbital. Dissected tissues were immediately fixed in 4% PBS-buffered formaldehyde and sent to the Mayo Clinic Histology Core Laboratory for subsequent sample processing and H&E and Masson's trichrome staining. Images were captured using the EVOS FL Auto Imaging System (ThermoFisher Scientific).

**RNA-binding protein immunoprecipitation**

RNA-binding protein immunuoprecipitation (RIP) assay was peformed using the EZ-Magna RIP kit (Sigma, 17-704) according to manufacturer's instructions. Briefly, 10 mg of right ventricle tissues from C57 mice were homogenized using a Bullet Blender (Next Advance). 1 ug of Anti-Sorbs2 antibody (Sigma, Cat#: SAB420183)) was used for the immunoprecipitation. Primers for quantitative RT-PCR to detect the Cx43 mRNA abundancy are Cx43/Gja1-F: 5'-ACAGCGGTTGAGTCAGCTTG-3', and Cx43/Gja1-R: 5'-GAGAGATGGGGAAGGACTTGT-3'.

**Participant recruitment**

From November 2010 to November 2017, 59 unrelated Han Chinese patients with ARVC were recruited with the approval from the Institutional Review Board of local hospital (IRB ID: TJ-C20181101). Specifically, 46 patients were recruited in Nanjing from the First Affiliated Hospital of Nanjing Medical University, and 13 patients were recruited in Wuhan from the Tongji Hospital,

Tongji Medical College, Huazhong University of Science and Technology. For controls, 402 gender- and ethnically-matched subjects without cardiomyopathy were recruited from a healthy population undergoing health examinations during the same period. All recruited individuals have undergone electrocardiogram, and echocardiography or computed (CT) or magnetic resonance imaging (MRI) tests to evaluated the structure and function of their hearts. Patients with ARVC were diagnosed based on the International Task Force Criteria of ARVC[8].

**Panel construction and targeted sequencing**

An ARVC panel was designed that can be used for ampliseq semiconductor next-generation sequencing (Ion Torrent, Thermo Fisher). The panel included 159.11kb full coding regions with more than 5 bp flanking regulatory sequences of 13 known genes for hereditary ARVC (Figure 6B in the online-only Data Supplement). The coding sequences of *CTNNA3* and *SORBS2* genes were later sequenced separately using Sanger sequencing. Primer sequences are listed in the Table 5 in the online-only Data Supplement). Genomic DNA was extracted from peripheral leukocytes using a standard protocol. Barcode-ligated libraries were constructed and mixed with the same molecular quantities. The pooled libraries were sequenced using an Ion Torrent sequencing platform.

**Whole exome sequencing (WES)**

In total, 200 ng of genomic DNA was fragmented into 250-350 bp fragments using sonication followed by end repairing. The DNA fragments were purified and ligated with adapters using SureSelect$^{XT}$ Library Prep Kit (Agilent, Santa Clara, CA). Next, the adapter-ligated DNA libraries were amplified and captured using the SureSelect capture library kit (Agilent, Santa Clara, CA). The captured sequences were further amplified for 350-bp read-length paired-end sequencing using Illumina X-ten system (Illumina, San Diego, California).

**Mutation identification and bioinformatics analysis**

The targeted sequencing raw data were initially processed with the Ion Torrent platform-specific software Torrent Suite v5.0. All reads were aligned to the GRCh37/hg19 human reference genome to analyze coverage status and to call variants. Extracted variants were annotated using Ion ReporterTM software 5.0 (Thermo Fisher). The WES data were processed according to the Genome Analysis Toolkit (GATK) Best Practices recommendations. All the potential pathogenic variants were identified with definite clinical significance according to recent American College of Medical Genetics (ACMG) guideline for the interpretation of sequence variants [9]. To identify potential pathogenic variants, we searched the Human Gene Mutation Database (HGMD) and the ClinVar database. All identified potential pathogenic variants that have been reported in HGMD or ClinVar databases were re-assessed whether linking to ARVC according to co-segregation data, identification of the variant in independent individuals, functional analyses of the variant. The expected phenotypes of these potential pathogenic variants should be consistent with carrier's clinical complications. Frequencies of all variants were searched from the 1000-Genome-Project, the ExAC, the ESP and the gnomAD databases. Missense variants were predicted using Polyphen-2, SIFT, and Mutation Taster. PhyloP score was used to predicted evolutionary conservation of variants. A variant was determined deleterious only when the variant was predicted to be damaging and conserved by all computational software. Having been validated via Sanger sequencing by using an Applied Biosystems 3500xl capillary sequencer (Applied Biosystems, Foster City) in order to eliminated false positive variants, each validated potential pathogenic variants was sequenced in 402 healthy controls using the same method. We then conducted Sanger sequencing of *SORBS2* gene in patients with ARVC. Finally, the variants were categorized into five classes including pathogenic, likely pathogenic, uncertain significance, likely benign, and benign.

**Minigene analysis**

Since human c.679+1G>T mutation is localized between intron 8 and intron 9 of *SORBS2* gene, a 2.4 kb-long DNA fragment was cloned from human *SORBS2* gene (NM_021069) covering part of the intron 9, whole exon 10, intron 10, exon 11, and part of the intron 11. The amplified DNA fragment was cloned into a Nde I site of pTBNde-min vector (Addgene) to express a wide type *SORBS2* fragment. To obtain the vector expressing a mutated SORBS2 fragment, mutagenesis was conducted using the site-directed mutagenesis system (Invitrogen) with primer pairs 5'-TCCTTATACATACAATGCAGTTAGGATGGGTTTCCTTGCTC-3' and 5'-GAGCAAGGAAACCCATCCTAACTGCATTGTATGTATAAGGA-3'. The vectors were transfected into AD-293 cell. 48 hour later , RNA were extracted from the transfected cell to check alternative splicing via RT-PCR, with 5'- CAACTTCAAGCTCCTAAGCCACTGC-3' as the forward primer, and 5'-TAGGATCCGGTCACCAGGAAGTTGGTTAAATCA-3' as the reverse primer.

**Zebrafish Morpholino injection and cardiac function quantification**

Zebrafish embryos were injected with *sorbs2a-e6* morpholino: 5'-TTTATTTGAAAGAACTCACCGTTGT-3' (0.25 μM), and/or *sorbs2b-e10* morpholino: 5'-ATGTGAAACAATAGGAAACCTTGTT-3' (0.25 μM) at one-cell stage. Heart fractional shortening (FS) and heart rate were measured at 3 dpf accordingly to our previously reported methods[10]. RT-PCR was conducted to assess knocking down efficiency of morpholinos. The exon skipping event in zebrafish injected with the *sorbs2a* morpholino was detected with the forward primer 5'-CCCAAGGACTGGTACAAGAC-3' and the reverse primer 5'-GAGGATTTTCCAGGCTCATACTC-3'. The intron retention event in zebrafish injected with the *sorbs2b* morpholino was detected with the forward primer 5'-GATTGGTACAAGACCATGTTCAAAC-3' and the reverse primer5'-GTCTGCACCCTTCTGCCAACAC-3'.
